# Reconstruction of dynamic regulatory networks reveals signaling-induced topology changes associated with germ layer specification

**DOI:** 10.1101/2021.05.06.443021

**Authors:** Emily Y. Su, Abby Spangler, Qin Bian, Jessica Y. Kasamoto, Patrick Cahan

**Author notes:** These authors made equal contributions (Emily Y. Su, Abby Spangler). Declarations of interest: none.

## Abstract

Elucidating regulatory relationships between transcription factors (TFs) and target genes is fundamental to understanding how cells control their identity and behavior. Computational gene regulatory network (GRN) reconstruction methods aim to map this control by inferring relationships from transcriptomic data. Unfortunately, existing methods are imprecise, may be computationally burdensome, and do not uncover how networks transition from one topology to another. Here we present Epoch, a computational network reconstruction tool that leverages single cell transcriptomics to infer dynamic network structures. Epoch performs favorably when benchmarked using data derived from *in vivo*, *in vitro,* and *in silico* sources. To illustrate the usefulness of Epoch, we applied it to identify the dynamic networks underpinning directed differentiation of mouse embryonic stem cells (ESC) guided by multiple primitive streak induction treatments. Our analysis demonstrates that modulating signaling pathways drives topological network changes that shape cell fate potential. We also find that Peg3 is a central contributor to the rewiring of the pluripotency network to favor mesoderm specification. By integrating signaling pathways with GRN structures, we traced how Wnt activation and PI3K suppression govern mesoderm and endoderm specification, respectively. Finally, we compare the networks established in *in vitro* directed differentiation of ESCs to those in *in vivo* gastrulation and mesoderm specification. The methods presented here are available in the R package Epoch, and provide a foundation for future work in understanding the biological implications of dynamic regulatory structures.

## Introduction

Gene regulatory networks (GRNs) model the regulatory relationship between a set of regulators, or transcription factors (TFs), and their target genes. The topology of these networks, defined by edges that map regulatory interactions between TFs and targets, offers a molecular-level view of a controlled system in which genes work together as part of a framework to accomplish specific cell functions (Karlebach and Shamir, 2008; Le Novère, 2015). Uncovering the topology of these GRNs is fundamental in answering a number of questions including: understanding how cellular identity is maintained and established (Davidson and Erwin, 2006), elucidating mechanisms of disease caused by dysfunctional gene regulation (Morgan et al., 2020; Qin et al., 2019), and finding novel drug targets among others (Carro et al., 2010). In the context of cell fate engineering, accurate mapping of GRNs would enable the identification of TFs required to activate expression of target genes of interest such that we could more accurately control cell fate transitions or cell behavior (Cahan et al., 2021; Rackham et al., 2016). Unfortunately, how best to map these relationships remains both an experimental and computational challenge.

Experimental approaches, including ChIP-ChIP and ChIP-seq, have provided valuable insights into identifying regulatory targets and TF binding site motifs of specific TFs in certain cell lines and cell types. Similarly, chromatin accessibility assays that detect transcription factor binding site (TFBS) footprints at regulatory genomic regions, such as ATAC-Seq (Buenrostro et al., 2013), enable the inference of regulatory networks. However, these types of experimental approaches are generally limited in scope (e.g. ATAC-defined networks are limited to TFs with known motifs) and scalability (e.g. it is infeasible to perform ChIP-seq for all TFs in all cell types). For these reasons, computational methods to efficiently infer GRN structures are needed. Existing methods have leveraged advances in data collection and machine learning approaches that have allowed for the statistical inference of GRNs via gene expression data. These include tools that rely on information theoretic measures (Faith et al., 2007; Margolin et al., 2006; Meyer et al., 2007), ensemble methods (Huynh-Thu et al., 2010), Bayesian approaches (Yu et al., 2004; Hartemink, 2005), or differential equation-based (ODE) approaches (di Bernardo et al., 2005). Unfortunately, computational methods developed to date suffer from several drawbacks including low precision and low sensitivity. Some of the leading contributors to the low performance include the difficulty in distinguishing direct from indirect interactions (Marbach et al., 2010), the confounding effects of Simpson’s Paradox (Trapnell, 2015), and the fact that bulk derived data does not offer perturbation sufficient to detect true regulatory relationships (Stark et al., 2003). Attempts to ameliorate these by aggregation across methods have met with modest success (Marbach et al., 2012).

With the advent of droplet-based systems for single-cell RNA-sequencing (Klein et al., 2015; Macosko et al., 2015; Zheng et al., 2017), and the corresponding rapidly growing body of single-cell transcriptomic data, computational analysis techniques have seen widespread advancement in harnessing the single-cell resolution to identify cell types and states (Villani et al., 2017) and to trace developmental trajectories (Treutlein et al., 2014). Of particular interest, trajectory inference (TI) methods, or pseudotemporal ordering methods, such as the transcriptomic-based methods Monocle (Trapnell et al., 2014), Slingshot (Street et al., 2018), and DPT (Haghverdi et al., 2016), as well as cytometry-based methods such as SPADE (Qiu et al., 2011), have allowed for computational modelling of dynamic processes like cell differentiation. Such methods order cells in an unbiased fashion along a trajectory following a linear, bifurcating, or more complex graph structure. In doing so, TI methods have proven valuable in the identification of novel cell subpopulations (Chen et al., 2016) and the elucidation of differentiation trees (Velten et al., 2017).

Unsurprisingly, several recently published tools have aimed to take advantage of the resolution offered by single-cell transcriptomics to infer GRN structures (Aibar et al., 2017; Matsumoto et al., 2017; Qiu et al., 2020). Just like their bulk counterparts, these methods can be categorized into information theoretic, ensemble, Bayesian, and ODE approaches. Unfortunately, they also suffer from similar limitations, and aside from low precision and sensitivity (Chen and Mar, 2018), many of these methods are computationally burdensome, limiting users’ ability to hone analysis through iterative application (Bonnaffoux et al., 2019). Additionally, though many of these methods employ pseudotemporal analysis to aid in reconstruction, they are limited in their ability to explain how network topology evolves over time. Importantly, we define a dynamic topology as one that exhibits changes in the edge-level regulatory relationships between TFs and targets over time (i.e. changes in the GRN connectivity) which thereby increases the number of reachable cell states (**Figure 1**). This implies that for a given TF-target relationship, the modulation of the TF in a particular context or time point would lead to changes in target expression (i.e. the relationship exists and is functional at one time point), but the modulation of the same TF in another context or time point would not lead to changes in target expression (i.e. the relationship is not functional at another time point). In other words, the dynamic and noncommutative nature of regulatory networks permits independent control of targets or genetic programs that would otherwise be simultaneously activated with non-sequential, or combinatorial, logic (Letsou and Cai, 2016). Such changes or shifts in network topology can arise for several reasons including the presence or absence of co-factors, the addition of epigenomic modifications, or changes in chromatin accessibility. Thus, for GRNs to more accurately model the emergence of distinct cell fates in development, they must encapsulate this dynamic behavior changing GRN topology. Moreover, because GRN topology dictates how a cell responds to perturbation, uncovering the dynamic GRN aids in understanding how the landscape of reachable cell states changes over time, which has significant implications in the quest to engineer cell fate.

**Figure 1.**
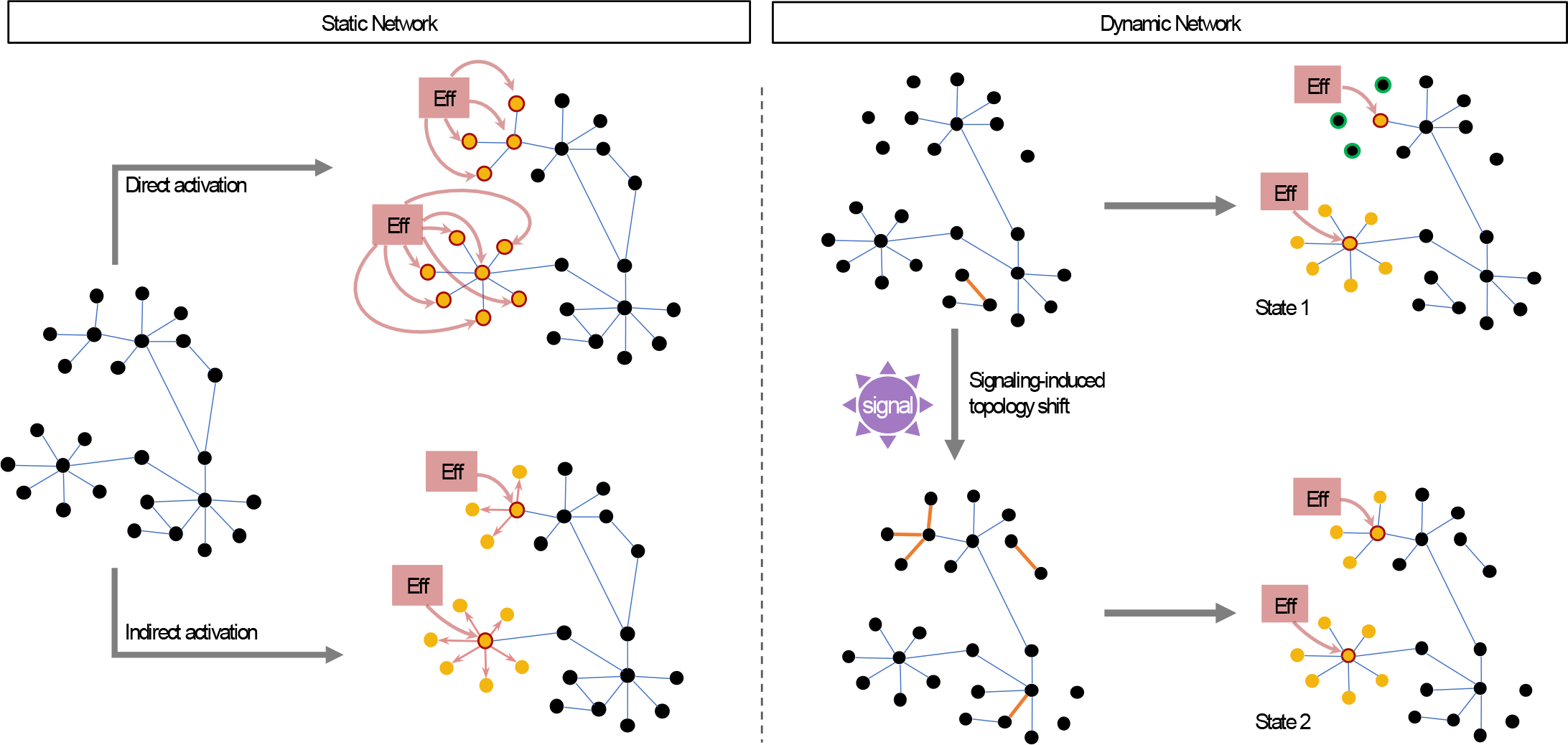
Effector activity on static and dynamic networks. Given a static GRN, manipulating signaling activity may result in an effector guiding cell state via two possibilities: directly, the effector may regulate a set of fate-specific genes, or indirectly, the effector may regulate a set of TFs that further regulate the same fate-specific genes. Both cases result in the activation of the same program. Alternatively, given a dynamic network, the state space of possible cell fates increases since the result of effector activity is dependent on a changing network topology. As a result, multiple sets of genetic programs may be regulated through the same signaling mechanism. Further, these shifts in GRN topology maybe be induced by the signaling activity itself.

For those reasons, we developed a computational GRN reconstruction tool called Epoch, which leverages single-cell transcriptomics to efficiently reconstruct dynamic network structures. Three key strategies exist at the center of Epoch: first, reconstruction is limited to dynamically expressed genes. Second, Epoch employs an optional “cross-weighting” strategy to reduce false positive interactions. Third, Epoch divides pseudotime into “epochs”, or time periods, and extracts a dynamic network and predicted top regulators influential in driving topology changes. Finally, Epoch includes a number of functionalities to aid in network analysis and comparison, including integration of the reconstructed GRN with major signaling pathways and the subsequent tracing through the GRN of shortest paths from signaling effectors toward activation or repression of target genes.

To demonstrate the utility of our tool we benchmarked Epoch’s performance against commonly used computational GRN reconstruction tools on synthetically generated data, *in vivo* mouse muscle development data, and *in vitro* directed differentiation data. We demonstrate that Epoch performs well in precision-recall analysis and in runtime comparison. We further highlight how Epoch’s stepwise-structured pipeline allows for user flexibility in adapting it to other GRN reconstruction methods.

Finally, we applied Epoch to answer open questions in gene regulation and directed differentiation. In particular, as pluripotent cells differentiate and undergo gastrulation, their transcriptomic profiles diverge, ultimately specifying and committing to distinct germ layers. Pluripotent cells undergoing *in vitro* differentiation often do so via embryoid body (EB) formation, which mimics *in vivo* gastrulation events such as primitive streak formation. Importantly, cell migration and specification *in vivo* are guided by a tightly-orchestrated combination of signals. Notably, Wnt3a, Bmp4, and Nodal concentration gradients (generated by Bmp and Wnt antagonist secretion on the anterior side of the embryo) are established early on, and are responsible for initiating primitive streak (Brennan et al., 2001; Yamamoto et al., 2004). For example, Wnt3a activates Brachyury in posterior epiblast cells, generating mesodermal cells (Perea-Gomez et al., 2002). Additionally, a number of FGFs (fibroblast growth factors) have roles in coordinating events in early embryo development including the exit from pluripotency. For example, primitive streak cells both synthesize and respond to FGFs, including Fgf4 and Fgf8, the absence of which results in failure of cells to migrate away from the primitive streak, resulting in the lack of mesodermal and endodermal tissue development and the disrupted patterning of the neuroectoderm (Sun et al., 1999). The orchestration of development by signaling pathways is mediated by the direct regulation of distinct transcriptional targets by pathway effectors (**Figure 1**). However, signaling cascades also have non-transcriptional consequences, including the alteration of epigenomic state and thus GRN topology (Fagnocchi et al., 2016; Mohammad and Baylin, 2010). The extent to which signaling pathways influence cell fate by altering GRN topology is unknown. To address this question, we performed scRNA-seq on day 0 through day 4 *in vitro* mouse embryonic stem cell (mESC) directed differentiation guided by four separate primitive streak induction treatments. We used Epoch to reconstruct GRNs associated with each treatment and compared the resulting topologies.

Specifically, we used Epoch to explore why cells treated with different induction treatments exhibited differing mesodermal fate potential, the lineage that was overall least likely chosen by differentiating cells. Our analysis revealed a number of topological features affecting mesodermal fate potential, including a central role for Peg3 in regulating the exit of pluripotency and transition to mesodermal fate. Additionally, by integrating signal transduction pathways with Epoch-reconstructed networks, we found that primitive streak induction via GSK inhibition produced a network topology that allows for more effective mesodermal specification. This was in direct contrast to other treatments in which network topologies more easily specified endodermal fate through suppression of PI3K signaling and subsequent activation of Foxo1. Indeed, there were remarkably large differences between target genes of the same TF across treatments, supporting a model where activation of signaling pathways directly alters network topology to influence cell fate potential. Finally, we reconstructed the dynamic GRN underpinning *in vivo* gastrulation and mesoderm specification for the purposes of comparison to our *in vitro* network. We identified *in vivo*-specific patterning modules that were not activated in the *in vitro* network, and further identified residual neuroectoderm modules present in the *in vitro* network. We believe that these results and similar analyses can be used to hone more efficient differentiation of mesodermal-fated cells. Further, we expect that analogous analyses can be performed to explain fate decisions in other dynamic biological processes.

## Results

### Epoch workflow relies on single-cell analysis techniques to infer dynamic network structures

Epoch takes as minimum input processed single-cell transcriptomic data and pseudotime or equivalent annotation from any trajectory inference method a user chooses to utilize (**Figure 2**). From here, Epoch employs its first step: limiting reconstruction to dynamically expressed genes in an effort to both focus on genes playing an active role in cell state changes, and to ensure that identified interactions are ones that can be inferred from perturbations in transcriptomic data. To accomplish this, Epoch models individual genes across pseudotime using a generalized additive model, and filters for significance. Alternatively, users have the option to employ TradeSeq (Van den Berge et al., 2020), a recently developed tool that employs a generalized additive model based on the negative binomial distribution to check for dynamic gene expression. Upon filtering the data for dynamically expressed genes, Epoch reconstructs an initial static network via a CLR-like (Context Likelihood of Relatedness) method, employing either Pearson correlation or mutual information to infer interactions (Faith et al., 2007).

**Figure 2.**
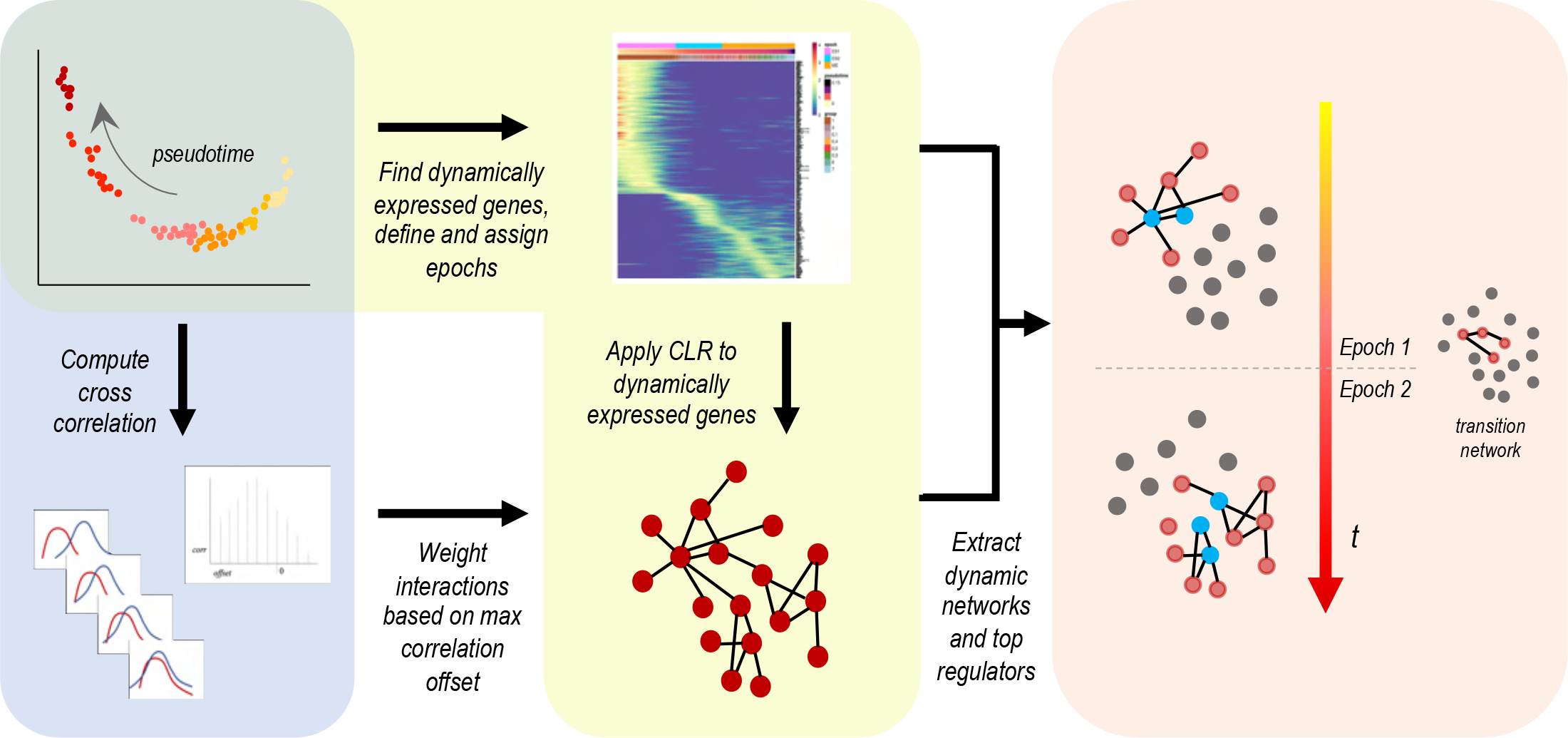
Epoch workflow. Epoch relies on a three-step process to reconstruct dynamic network structures: (1) extraction of dynamic genes and CLR, (2) application of cross-weighting, and (3) extraction of the dynamic network.

In the second step, which we have termed “cross-weighting”, Epoch employs an optional network refinement step that relies on a cross-correlation-based weighting scheme. This helps to reduce false positives that may result from indirect interactions or that may represent non-logical interactions, ultimately improving precision (**Supplementary Figure 1a**). In this step, the expression profiles over pseudotime for each TF-target pair is aligned and progressively shifted to determine an average estimated offset value at which maximum correlation is achieved between the two profiles. A graded-decline weighting factor is computed based on this offset, and used to negatively weight interactions that are less likely to be true positives.

Once a static network has been inferred, Epoch extracts a dynamic network from the static network, and further identifies the top regulators driving changes in network topology and thus cell state. Specifically, this begins with Epoch breaking down pseudotime into “epochs”, or discrete time periods, based on pseudotime, cell ordering, k-means or hierarchical clustering, sliding window similarity, or user-defined assignment. With the exception of user-defined assignment, all methods of defining epochs are, to some extent, automated, with epoch definitions learned directly from the data (see Methods). From here, genes are assigned to epochs based on their activity along pseudotime (a given gene may be assigned to multiple epochs). Epoch then fractures the static network into a dynamic one, composed of “epoch networks”, representing active interactions within a particular epoch, and possible “transition networks”, describing how an epoch network transitions into a subsequent epoch network.

As Epoch was initially built with the improvement of cell fate engineering protocols in mind, we included in the Epoch framework a number of network analysis and network comparison capabilities. For example, Epoch will identify influential transcription factors within a given static or dynamic network by extracting top regulators that appear to have the most influence on network state. Specifically, Epoch will compute and rank TFs according to their PageRank (Brin and Page, 1998) in the context of their epoch networks, and will further look for TFs that simultaneously score high in betweenness and degree centralities as compared to all other TFs. Additionally, Epoch can integrate major signal transduction pathways with reconstructed networks. This can be used to determine the shortest paths through the networks from pathway effectors to functional groups of specified target genes. This allows users to identify and trace topologies capable of activating or repressing specific groups of genes.

Finally, Epoch is modularly designed such that it can be broken into individual steps and flexibly merged and used with any trajectory inference, network reconstruction, and even network refinement tools that currently exist or that may be developed in the future.

### Epoch performs favorably on synthetically generated data

To assess the performance of Epoch in reconstructing static networks, we benchmarked it on synthetically generated data. We generated fifteen synthetic datasets using the Dyngen R package (Cannoodt et al., 2020). Networks were designed to encompass varying regulatory motifs and spanned 100-420 genes including 20-70 transcription factors. Synthetic single-cell datasets generated from these networks varied in size, from as low as 20 to as high as 1000 cells. We compared Epoch to the original versions of GENIE3 (Huynh-Thu et al., 2010), which is the GRN reconstruction engine of SCENIC (Aibar et al., 2017), and CLR (Faith et al., 2007) as well as variations on these methods that incorporated them within the Epoch framework. Thus, we assessed a total of 11 methods: CLR using mutual information (original CLR), CLR using Pearson correlation, CLR using mutual information with cross-weighting, CLR using Pearson correlation with cross-weighting, GENIE3 (original GENIE3), GENIE3 limited to dynamically expressed genes, GENIE3 limited to dynamically expressed genes with cross-weighting, Epoch with Pearson correlation (with and without cross-weighting), and Epoch with mutual information (with and without cross-weighting).

We reconstructed individual networks with each method for each dataset, using normalized simulation time as the pseudotemporal annotation for Epoch. To assess performance, we compared each resulting network to a set of five random networks, unique to each reconstructed network, that were generated by permuting edge weights of targets for each transcription factor in the reconstructed network (**Figure 3a**). As a gold standard, all networks were compared against the original network used to generate the synthetic single-cell data. Based on fold improvement in area under the precision-recall curve over the random networks, all variations of Epoch outperformed original versions of CLR and GENIE3 (**Figure 3b, Supplementary Figure 1b**). Additionally, we found that Epoch using mutual information with or without cross-weighting resulted in higher fold improvement over random compared to any other method. The addition of cross-weighting improved median AUPR fold change over random for all methods. Further, limitation to dynamic genes (e.g. versions of CLR vs. corresponding versions of Epoch as well as GENIE3 vs. GENIE3 limited to dynamic genes) also improved performance. In all cases, it was apparent that key steps in the Epoch framework improved overall static reconstruction, as limiting reconstruction to dynamically expressed genes and applying the cross-weighting scheme both led to increased fold improvement in AUPR over random, demonstrating the utility of Epoch’s framework.

**Figure 3.**
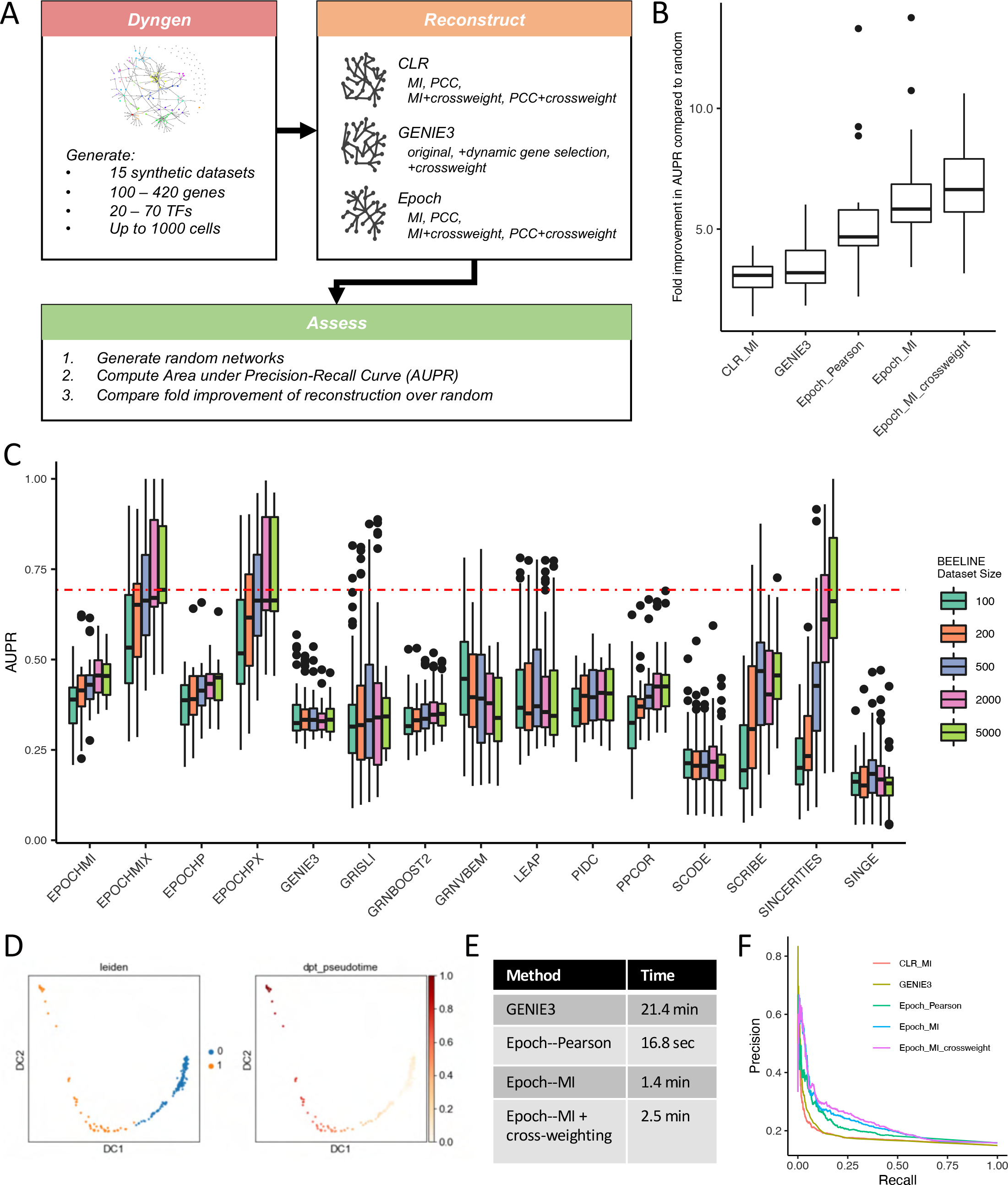
Benchmarking Epoch. (A) Synthetic dataset generation via Dyngen, reconstruction, and assessment. (B) Fold improvement in AUPR over random of three versions of Epoch (Pearson, MI, and MI with cross-weighting) against CLR and GENIE3 on GRN reconstruction from Dyngen-generated synthetic data; Kruskal-Wallis p=4.26×10^-8^. (C) BEELINE benchmarking on synthetic data. Epoch using MI and Pearson denoted with “MI” and “P” respectively. “X” indicates use of cross-weighting. Red line indicates the median AUPR of the best performing method, Epoch using MI+cross-weighting on size 5000 cell datasets. Kruskal-Wallis p<2.2×10^-16^. (D) Clustering and diffusion pseudotime for E12.5 muscle development data used in benchmarking. (E) Runtimes for reconstruction on muscle development data. (Runtimes were measured on a MacBook Pro laptop with 2.9 GHz Intel Core i7 processor, 16GB 2133 MHz DDR3 of RAM). (F) Precision-recall curves for three versions of Epoch (Pearson, MI, and MI with cross-weighting), CLR, and GENIE3 based on reconstruction from muscle development dataset.

We next compared Epoch to recently developed GRN reconstruction methods designed for single-cell data. We used the benchmarking platform, BEELINE (Pratapa et al., 2020), to assess Epoch’s performance against 11 other single-cell methods across the synthetic datasets available within the tool (**Figure 3c, Supplementary Figure 2**). Importantly, these covered a range of cell state trajectories (linear, long linear, bifurcating, bifurcating converging, cycle) as well as dataset sizes (100 to 5000 cells). Our results demonstrate that Epoch with cross-weighting (using either mutual information or Pearson correlation) performed best based on AUPR. Additionally, while SINCERITIES almost approaches Epoch’s performance at the 5000- cell dataset size, it performs markedly worse with smaller datasets, unlike Epoch. Finally, additional comparison of computational runtimes demonstrated that Epoch has the additional benefit of running magnitudes more efficiently than other methods including GRNBOOST2, PIDC, and GENIE3 (**Supplementary Figure 3**).

We then compared Epoch against the same 11 single-cell GRN reconstruction methods across the curated datasets available in BEELINE (**Supplementary Figure 4**). Importantly these datasets, though synthetic, were simulated from literature-derived networks and pseudotime was computed using Slingshot (Street et al., 2018). In this case performance based on AUPR was more equal across the methods, with GENIE3, GRNBoost2, and PIDC slightly out-performing other methods (**Supplementary Figure 4a**). On closer inspection, it was clear that performance was dataset dependent. For example, Epoch and other pseudotime-based methods performed relatively better in reconstructing the GSD network vs the HSC network (**Supplementary Figure 4b,c**). The same phenomenon was present in the BEELINE publication where it was initially attributed to imprecision of trajectory inference in estimating pseudotime. To test this, we used the underlying literature derived-networks and re-simulated the datasets. We then used the true simulation time, rather than Slingshot-derived pseudotime, for reconstruction and benchmarking. Our results demonstrated that despite using the true simulation time, AUPR of pseudotime-based reconstruction methods was largely unchanged, suggesting that poor pseudotime computation was not at fault (**Supplementary Figure 4d**).

Finally, we hypothesized that decreased performance in the pseudotime-based methods was largely driven by reconstruction on split trajectories. Namely, for pseudotime-based methods, reconstruction of branched data was performed along each lineage separately and resulting networks were aggregated. For example, HSC datasets represented four distinct lineages (branches), and so the 2000 cell dataset was broken into roughly four 500 cell datasets for reconstruction by pseudotime-based methods before aggregation. For non-pseudotime-based methods, including GENIE3, GRNBoost2, and PIDC, reconstruction was done on the full 2000 cell dataset. To test the extent to which this impacted performance, we re-benchmarked GENIE3, GRNBoost2, and PIDC splitting the reconstruction along each lineage before aggregating, mirroring the pseudotime-based methods. The resulting performance was substantially decreased, suggesting that splitting reconstruction is a factor contributing to decreased performance (**Supplementary Figure 4e**). Ultimately, other factors, such as network density and the presence of specific connectivity motifs, warrant further exploration, but was unfortunately infeasible with the limited data.

### Epoch performs favorably on real world data

While the results from assessment of networks reconstructed from synthetic data demonstrated Epoch’s favorable performance as a network inference tool, we acknowledge that there are caveats associated with synthetic data, including limited stochasticity and reliance on simulations that may not encompass the full extent of potential cell state responses seen in real biological systems. Thus, we next turned our attention to benchmarking on real world data. To assess Epoch’s performance, we began by assessing the networks reconstructed by the variations of Epoch, CLR and GENIE3 listed above from data of E12.5 mouse muscle development (**Figure 3d**). For this dataset, we used Diffusion based pseudotime (DPT) to compute pseudotime for Epoch reconstruction (Haghverdi et al., 2016). We used Chip-X data from the Enrichr database, which includes ChIP-Seq, ChIP-ChIP, ChIP-PET, DamID used to profile transcription factor binding to DNA, covering 220 TFs and >35,000 genes, as a gold standard network (Lachmann et al., 2010). Based on precision recall, we found that all variations of Epoch once again outperformed original versions of CLR and GENIE3. Interestingly, we found that the two variations of GENIE3 embedded within the Epoch framework performed best overall in this assessment, followed by the two variations of Epoch using mutual information (with and without cross-weighting). It was again apparent that limiting reconstruction to dynamically expressed genes and cross-weighting significantly improved overall reconstruction (**Figure 3f, Supplementary Figure 1c**).

To determine if these results were consistent across other real world datasets, we turned to single-cell transcriptomic data collected from day 4 in vitro mESC directed differentiation (Spangler et al., 2018). We limited our reconstruction to cells in earlier stages of differentiation that had not yet transitioned into any committed germ layer-like state, as determined by pseudotemporal analysis, RNA velocity analysis, expression of Zfp42, marking naive pluripotent cells, and expression of Fgf5, marking cells in a primed state (**Supplementary Figure 1d**). The filtering ensured the trajectory remained linear, which, while absolutely not a requirement for Epoch reconstruction, was employed to minimize performance variability of trajectory inference methods in which branched structures may be more difficult to order. This allowed us to narrow our focus on reconstruction performance while minimizing impact from potentially inaccurate pseudotemporal ordering. Based on precision-recall, we found that Epoch variations using cross-weighting performed best, followed by GENIE3 embedded within the Epoch framework (**Supplementary Figure 1e**). Overall, these benchmarking results demonstrate Epoch’s utility as a reconstruction tool, and further demonstrate the usefulness of Epoch as a flexible framework that can improve reconstruction of other methods.

### Single cell RNA-sequencing of early in vitro mouse ESC directed differentiation

With the goal of exploring the dynamic GRN topology underlying lineage specification in gastrulation, we collected scRNA-seq data from day 0 through day 4 *in vitro* mESC directed differentiation guided by four separate treatments encouraging primitive streak formation (**Figure 4a**). Briefly, cells were allowed to differentiate in serum-free differentiation media before being treated with one of four treatments on day 2: Wnt3a and Activin A alone (WA), or with one of Bmp4 (WAB), GSK inhibitor (WAG), or Noggin (WAN). Multiple rounds of differentiation were staggered such that we could harvest samples representative of d0-d4 at one time for sequencing using the MULTI-seq protocol (McGinnis et al., 2019). After barcode classification, samples were preprocessed using the SCANPY pipeline (Wolf et al., 2018), and RNA velocity analysis was performed (Bergen et al., 2019; La Manno et al., 2018) (**Figure 4b,c**).

**Figure 4.**
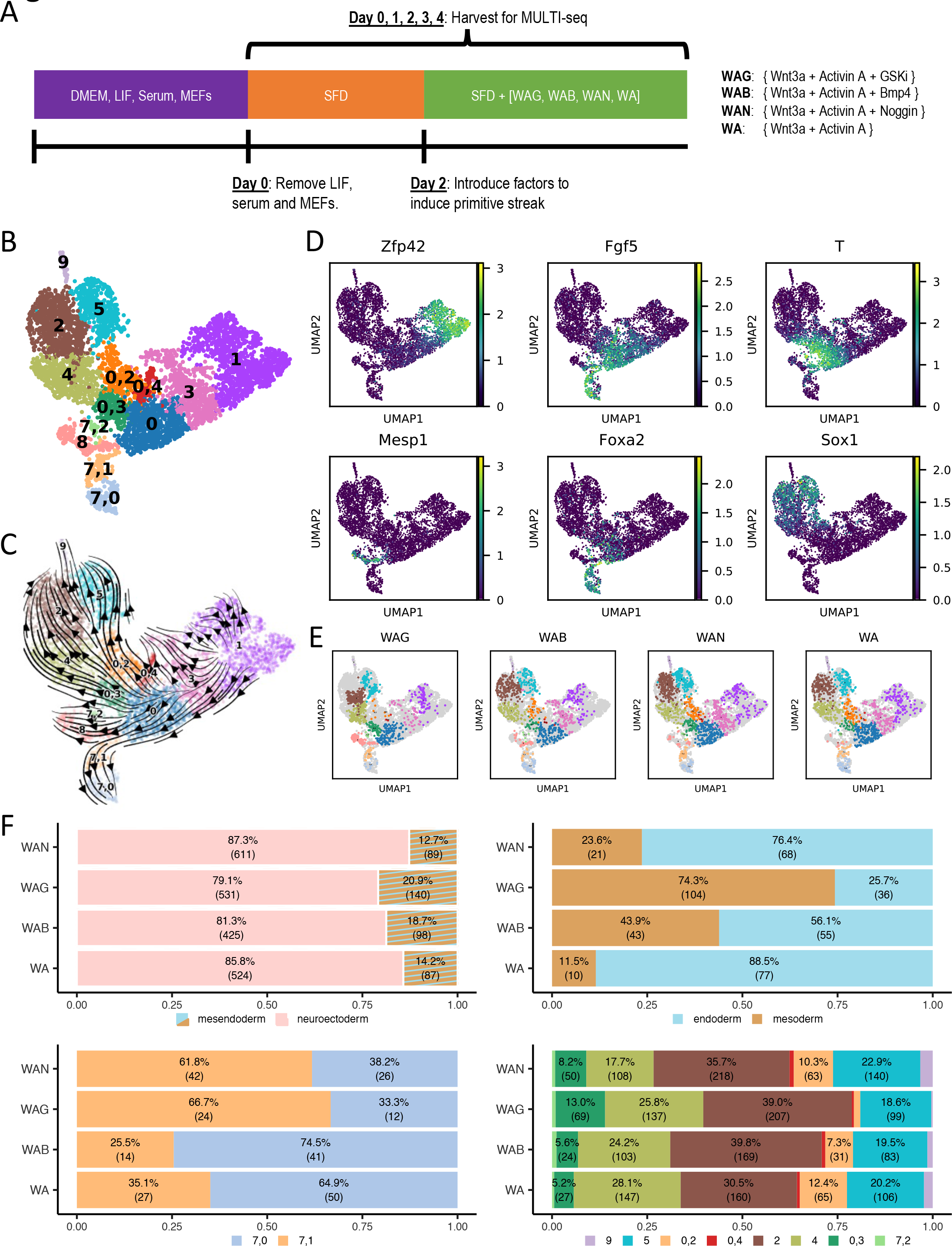
mESC in vitro directed differentiation. (A) Directed differentiation protocol. Cells were harvested from d0 through d4 samples for MULTI-seq. (B) Clustering for MULTI-seq data. (C) RNA velocity for MULTI-seq data. (D) Select marker gene expression. (E) Cell populations by treatment. (F) Quantification of cell fate distribution based on treatment comparing: mesendoderm vs. neuroectoderm, endoderm vs. mesoderm, cluster 7,0 vs. 7,1 along endoderm fate, and cluster distribution along neuroectoderm fate.

We found the presence of three distinct lineages, which we confirmed to be representative of neuroectoderm (lineage expressing Sox1), mesoderm (lineage expressing Mesp1), and endoderm (lineage expressing Foxa2) based on gene expression and further verified by singleCellNet (SCN) analysis (Tan and Cahan, 2019) (**Figure 4d, Supplementary Figure 5, 6a**). RNA velocity additionally supported our cluster annotations. We found that the majority of the differentiating cells transitioned toward the neuroectoderm (clusters labeled 0,2, 0,3, 4, 2, 5, 9) fate with smaller populations transitioning toward mesoderm (cluster labeled 8) and endoderm fate (clusters labeled 7,1 and 7,0).

Undifferentiated cells populated cluster 1, which was positive for Zfp42 expression, a pluripotency marker (Rogers et al., 1991). This population additionally classified as inner cell mass, further supporting their pluripotent annotation. RNA velocity indicated these cells transitioned into cluster 3, which marked the beginning of Fgf5 expression, an epiblast maker (Haub and Goldfarb, 1991; Pelton et al., 2002). SCN revealed early ectoderm and epiblast signature for cluster 3, indicating to us that it was representative of a primed cell state. Cells in this cluster transitioned to cluster 0 and 0,4. Cells in cluster 0 were largely representative of primitive streak, as determined by Brachyury expression, with some early ectoderm and residual epiblast signature, as determined by SCN. Interestingly, cluster 0,4, which expressed a lower level of Brachyury and exhibited some primitive streak and early ectoderm SCN signature, was comprised of day 0 cells. It is possible that these cells, which had yet to undergo the differentiation protocol, had escaped LIF-controlled naïve pluripotency and were already in a primed early ectodermal state.

Cluster 0 cells expressing Mixl1, a regulator of mesoderm and endoderm (Hart et al., 2002) further branched into two distinct lineages according to RNA velocity: mesoderm (cluster 8) and endoderm (clusters 7,1 and 7,0). Cluster 8 was largely enriched for Mesp1, and some cells at later time (determined by RNA velocity) in this cluster also expressed Tbx6. SCN analysis revealed that this cluster exhibited multiple mesoderm-related signatures including primitive streak, anterior paraxial mesoderm, presomitic mesoderm, and somites, ultimately supporting its mesoderm annotation. Along the endoderm path, cluster 7,1 transitioned from cluster 0 and further transitioned into cluster 7,0. Both 7,1 and 7,0 are enriched for Foxa2 and Sox17 expression. SCN analysis suggested an increasing endodermal signature down this path, with cluster 7,1 classifying as anterior primitive streak and weaker as definitive endoderm and gut endoderm, while cluster 7,0 more strongly classified as definitive and gut endoderm.

The neuroectodermal path begins from both cluster 0 and 0,4 according to RNA velocity results. We annotated clusters 0,3 and 4 as neuromesodermal progenitor (NMP) populations. SCN further verified cluster 0,3, which expresses T and Sox2, as early NMP and cluster 4, which expresses Sox2 and exhibits variable T expression, as late NMP. Cluster 7,2, which expresses Chordin, classified strongly as node. Interestingly, RNA velocity suggested the existence of two parallel paths along the neuroectodermal lineage, with cluster 0,2 transitioning into cluster 5, and cluster 4 transitioning into cluster 2. Along the first path, cluster 0,2, classified weakly as a number of ectodermal cell types including early ectoderm. Because RNA velocity indicated this cluster transitioned into the later neuroectoderm cluster 5, we annotated it as early ectoderm. Cluster 5 classified strongly as forebrain/midbrain, with some weaker future spinal cord classification. Given this, and it’s Zic1 and En2 expression, we annotated this cluster as future brain. Cluster 9, which expresses Crabp1 and differentially expresses Tubb3 (suggesting a later neuronal identity compared to cluster 2 or 5), seems to transition from this cluster, and is thus annotated as future brain. Along the second neuroectodermal path, cells in cluster 2 classified more strongly as future spinal cord with a weaker forebrain/midbrain classification. Importantly this cluster expresses many patterning genes from the Hox family indicative of spinal cord fated cells, including Hoxd9 and Hoxb6 (Philippidou and Dasen, 2013). For this reason, we annotated this cluster as future spinal cord. A summary of our cluster labels can be found in the supplementary materials (**Supplementary Table 1**).

Interestingly, in annotating the data, it is apparent that there are differences between induction treatments in terms of the distribution of cells amongst the different populations (**Figure 4e,f, Supplementary Figure 6b**). Notably, cells treated with WAG exhibited a stunted trajectory toward neuroectodermal fate but a fuller trajectory toward mesodermal fate. This was in direct contrast to cells treated with the remaining three treatments (WAB, WAN, WA), in which very few cells differentiated toward mesodermal fate, but instead exhibited strong neuroectodermal or endodermal commitment. Further, while WAB and WA cells tended to transition into cluster 7,0 (furthest along endodermal development), WAN and WAG cells were more likely to stay in cluster 7,1.

### Epoch elucidates regulatory networks driving lineage specification in early in vitro mouse ESC differentiation

To begin our network analysis exploring lineage specification, we applied Epoch to the day 0 through day 4 directed differentiation data, using latent time (Bergen et al., 2020) to order cells, and reconstructing the networks underlying each of the three lineages. Latent time was divided into three epochs, and a dynamic network was extracted for each lineage (**Figure 5, Supplementary Figure 7**). In examining the Epoch-predicted top regulators and their top targets, we observed noteworthy regulatory programs emerging over time along the three lineages. For example, a module including the genes Gsc, Lhx1, Sox17, Foxa1, Foxa2, and Gata6 emerges in the third epoch along the endoderm path. Sox17 and Foxa2 serve as endoderm markers, as knockout of Sox17 has been shown to deplete definitive gut endoderm and lack of Foxa2 results in failure to develop fore- and midgut endoderm (Dufort et al., 1998; Kanai-Azuma et al., 2002). Foxa1 has also been implicated in the initiation and maintenance of the endodermal lineage (Ang et al., 1993), while inactivation of Lhx1, which has been shown to bind to the enhancer region of Foxa2, disrupts the development of definitive endoderm (Costello et al., 2015). The regulator Sox11 and a collection of Hox genes emerge in the third epoch of the neuroectoderm lineage, suggesting the activation of programs that pattern brain and spinal cord (Philippidou and Dasen, 2013). Ultimately, these networks served as a foundation for the remainder of the analyses in this study, representing TF-target regulation present along each path of lineage specification.

**Figure 5.**
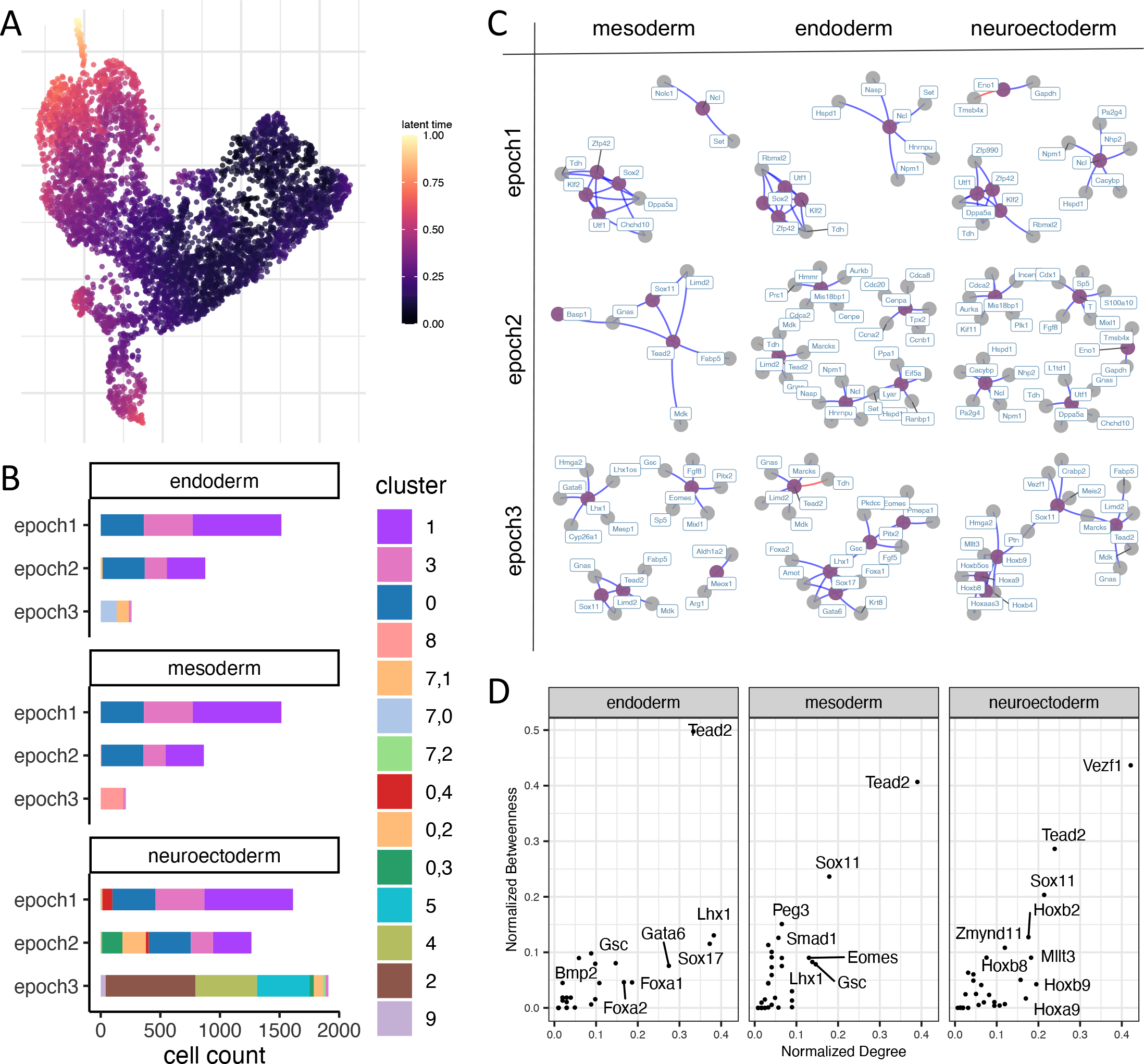
MULTI-seq directed differentiation network reconstruction. (A) Latent time annotation for MULTI-seq data. (B) Population breakdown by cluster of epoch assignment along each germ layer path. (C) Reconstructed dynamic networks along mesoderm, endoderm, and neuroectoderm paths, with top 5 regulators colored purple and their top targets in gray as determined by PageRank. (D) Top regulators as predicted by Epoch via betweenness and degree for the third epochs along each germ layer path.

In addition to directly extracting top regulators via PageRank and Betweenness-Degree from the reconstructed lineage networks, we highlight a more nuanced methodology to resolve influential TFs based on bagging, since the consensus of many reconstructed networks would be less susceptible to noise. Specifically, we limited our scope to early time, corresponding to a naïve to primed pluripotency transition, and bootstrapped the network reconstruction (**Supplementary Figure 8**). For each of these sampled and reconstructed networks, we predicted the top 10 regulators involved in the transition by both PageRank and Betweenness-Degree. We then applied a consensus approach in which regulators were ranked by their frequency as a top 10 regulator (based on PageRank) or average importance (based on Betweenness-Degree) amongst the bootstrapped networks. The top regulators recovered by both PageRank and Betweenness-Degree were largely overlapping, and included TFs such as Zfp42, Klf2, Sox11, Pou3f1, and Klf4.

Of the top 10 regulators predicted through this consensus approach, Klf2 and Klf4 are critical in maintaining ground state pluripotency (Yeo et al., 2014). For example, it has been demonstrated that nuclear export of Klf4 causes rapid decline in Klf4 transcription, a trigger that leads to the exit of naïve pluripotency, and the blocking of which prevents ESC differentiation (Dhaliwal et al., 2018). Thus, it is unsurprising that they are predicted to have an influential role in dictating network topology. Interestingly, a number of predicted top regulators, such as Pou3f1 and Sox11, are known to promote neural fate (Barral et al., 2019; Wang et al., 2013; Zhu et al., 2014). Knockdown of another predicted top regulator, Tcea3, has also been shown to bias differentiation toward mesendodermal fates (Park et al., 2013). Taken together, this suggests that the underlying network is primed for the neuroectodermal specification fate choice, aligning with the default model of neural induction in mESCs (Muñoz-Sanjuán and Brivanlou, 2002; Tropepe et al., 2001) and the apparent preference for neuroectoderm fate amongst cells in this dataset.

### Peg3 is a central regulator in mesodermal WAG networks and may drive differences in gene module expression amongst treatments

We next focused our attention on mesodermal fate specification in mESC directed differentiation. Specifically, in observing both the scarcity of the mesodermal population (in comparison to neuroectodermal fate) as well as the discrepancy amongst treatments in propensity for reaching mesodermal fate, we asked why WAG treated cells had the greatest potential for reaching this population. We hypothesized that underlying these differences in fate potential amongst treatments were treatment-specific GRNs. To this end, we reconstructed and compared dynamic networks for each treatment along the mesodermal path (which we refer to as the treatment-specific networks: WAG, WAB, WAN, and WA networks), and further compared these against the full reconstructed mesoderm network described in the previous section (which is reconstructed from all cells regardless of treatment, and which we refer to as the mesoderm network). We then aimed to understand why WAG treated cells were able to specify mesodermal fate to a greater extent than WAB, WAN, and WA treated cells.

We then sought to understand what regulators could be driving differences in mesoderm reachability between the treatments. We expected treatment-driven differences in fate potential to logically arise after cells undergo treatment on day 2, and would be most apparent toward later latent time. We also reasoned that we could quantify and monitor differences in cell state based on the activity of gene modules, defined as a community or unit of nodes that are highly interconnected, within the network. Specifically, understanding the activity of these modules in each treatment group would elucidate the extent to which the mesoderm state is established and further aid in identifying regulators that fail to activate these regulatory programs. Thus, we performed community detection on epoch 2, epoch 3, and their transition, of the mesoderm network and identified distinct TF communities, or modules, within the topology (**Figure 6a, Supplementary Figure 10**). We assessed the activity of each module by looking at the average expression of member genes across latent time for each treatment (members predicted to be repressed were not included in the average so as not to improperly depress this measure of activity) (**Figure 6b, Supplementary Figure 11a**). Importantly, we found the existence of certain modules that were strongly activated in the WAG treatment that failed to activate or were weakly activated in the remaining three treatments. Three such modules (one activated in each of the second epoch, third epoch, and transition) were particularly interesting as they exhibited strong activity in the WAG treatment along not only the mesoderm path, but also the full dataset. The TFs in these modules were Peg3, Lhx1, Hoxb2, Foxc1, Tshz1, Meis2, Mesp1, Tbx6, Foxc2, Prrx2, Meox1, and Notch1. Upon extracting the differential network (which we define as the network containing the interactions, or edges, that have large weight differences between two treatments, and thus represent interactions most likely to be unique to a treatment network; see methods) between WAG and WA, we recovered the same modules and further identified the top differential regulators of these TFs themselves (**Figure 6c, Supplementary Figure 11c**). Interestingly, in contrast to the WAG treatment, Peg3 was predicted to have a negligible role in the WA treatment.

**Figure 6.**
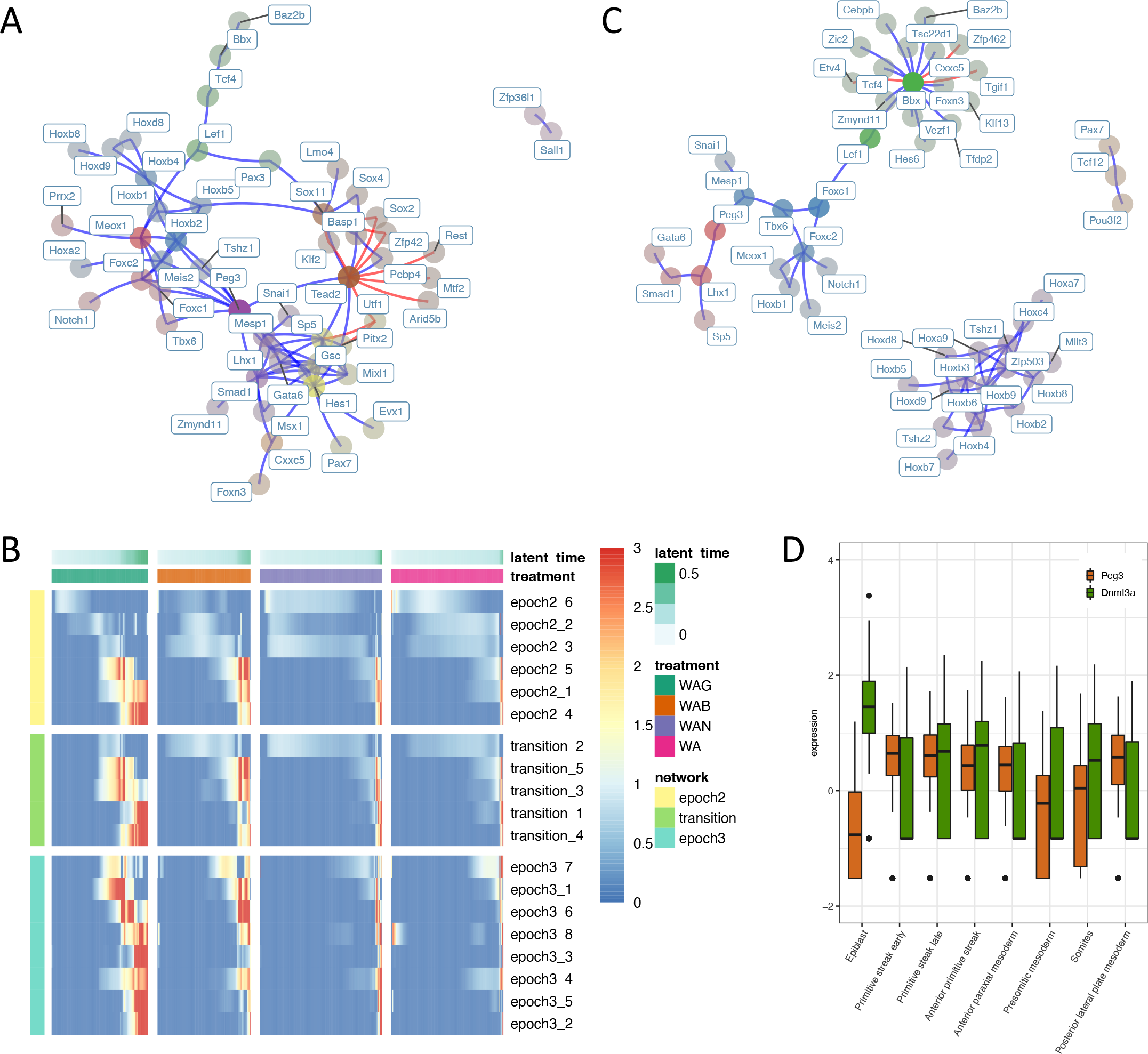
Mesodermal network analysis. (A) The epoch 3 subnetwork of the mesodermal dynamic network. TFs are colored by community and faded by betweenness. Blue and red edges represent activating and repressive edges respectively. (B) Average community expression over time by treatment along the mesodermal path. Communities shown (each row) are from the epoch 2 subnetwork (yellow), transition (green), and epoch 3 subnetworks (aqua). (C) The epoch 3 differential network between WAG (on) and WA (off). Interactions in this network represent edges present in the WAG mesodermal network that are not present in the WA mesodermal network. TFs are colored by community and faded by betweenness. Blue and red edges represent activating and repressive edges respectively. (D) Peg3 and Dnmt3a expression in a sampled portion of relevant cell types in gastrulation data from Grosswendt et al. Peg3 ANOVA p<2×10^-16^, Dnmt3a ANOVA p<2×10^-16^.

Peg3’s central influence in the WAG network, however, implied a pathway linking GSK3 inhibition, Peg3 expression, and mesodermal gene expression. Indeed, previous reports have demonstrated a role for GSK3 in maintaining DNA methylation at imprinted loci, including Peg3, in mESCs (Meredith et al., 2015). Additional literature has demonstrated that methylation at imprinted loci may be maintained by Dnmt3a2, which has been suggested to be regulated by the GSK3-dependent transcription factor N-Myc (Popkie et al., 2010). Unfortunately, our data was only able to capture Dnmt3a dynamics (representing composite Dnmt3a and Dnmt3a2 expression), and not specifically Dnmt3a2, which is transcribed from an alternative promoter within the Dnmt3a locus. However, since Dnmt3a2 is the predominant isoform in ESCs (Chen et al., 2002), we roughly compared Dnmt3a temporal dynamics and Peg3 expression between the different treatments (**Supplementary Figure 11b**). Notably, Dnmt3a and Peg3 expression were largely mutually exclusive, and there was low-to-no Dnmt3a expression in the WAG treated cells. We then asked if this phenomenon occurs in *in vivo* gastrulation. In the single-cell mouse gastrulation dataset from Grosswendt et al. (Grosswendt et al., 2020), we found relatively high expression of Dnmt3a in epiblast cells which was tapered in primitive streak and mesodermal cells (**Figure 6d**). In contrast, Peg3 expression was low in epiblast and increased in the primitive streak, mirroring our *in vitro* data, and suggesting that Peg3 may have a role in orchestrating the exit of pluripotency and specification of mesodermal fate *in vivo*.

### Tracing signaling cascades to germ layer transcriptional programs

We next sought to trace pathways from signaling effectors to the activation of mesodermal fate. To determine which signal transduction pathways were active at any given point in latent time along the mesodermal lineage, we used ChIP-seq data from ChIP-Atlas (Oki et al., 2018) to compile a list of binding targets for 18 signaling effector TFs (available as part of the Epoch framework). We then computed average expression of these targets across latent time broken down by treatment, and unsurprisingly found differences in the signaling pathways that were activated in each treatment (**Figure 7a**). Of note, Notch signaling was activated in WAB, WAN, and WA treatments, but not in the WAG treatment along the mesodermal lineage (and weakly activated along the neuroectodermal lineage), consistent with literature demonstrating that Notch signaling plays a role in specifying neuroectodermal fate in the neuroectoderm-mesendoderm fate decision (Androutsellis-Theotokis et al., 2006; Aubert et al., 2002).

**Figure 7.**
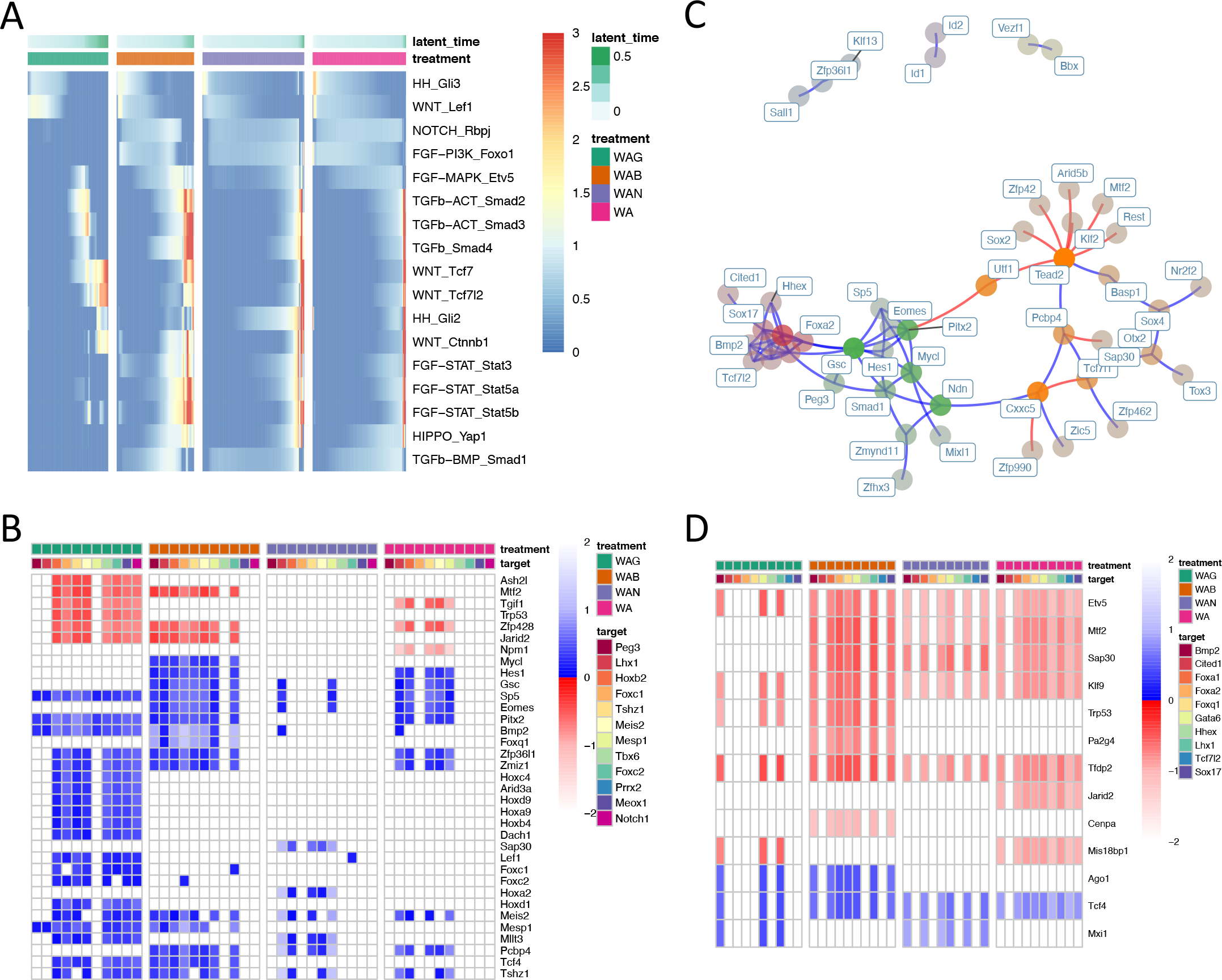
Integrating signaling effectors and tracing paths to target genes. (A) Average expression of targets of signaling effectors by treatment over time. Pathway and effector are labeled on each row. (B) Shortest path analysis showing length of shortest paths from Wnt effector targets (each row) to mesodermal target genes (each column) within treatment-specific networks. Path lengths are normalized against average path length within a network. Blue and red indicate paths that activate and repress the target gene respectively. If such a path does not exist, it has length of infinity and is therefore white. (C) The epoch 3 subnetwork of the endodermal dynamic network. TFs are colored by community and faded by betweenness. Blue and red edges represent activating and repressive edges respectively. (D) Shortest path analysis showing length of shortest paths from PI3K suppression (targets of Foxo1 activation) (each row) to endodermal genes (each column) within treatment-specific networks. As before, path lengths are normalized against average path length within a network. Blue and red indicate activation and repression of the target gene respectively. If a path does not exist, it has length of infinity and is therefore white.

Because Wnt signaling is essential for mesodermal fate, we looked for paths connecting Wnt effectors to the TFs in the WAG-specific modules defined previously. Specifically, we computed the shortest paths from targets of these effectors to the module TFs in each treatment-specific dynamic network (using edge lengths inversely proportional to the cross-weighted score) (**Figure 7b**). Paths were assigned infinite length if such a path did not exist. We normalized these path lengths against the average path length in a given network, and found that for the WAB, WAN, and WA treatments, no paths from Wnt to many of the module TFs existed.

Our signaling effector target analysis also revealed increased Foxo1 activity in WAB, WAN, and WA treated cells as compared to WAG treated cells, indicating the suppression of PI3K signaling along the WAB, WAN, and WA trajectories (**Figure 7a**). Importantly, it is known that the establishment of definitive endoderm (DE) requires suppression of PI3K signaling (McLean et al., 2007; Yu et al., 2015). Further, it has been recently shown in hiPSC to DE differentiation that Foxo1 binds to DE-formation-related genes and that inhibition of Foxo1 during DE induction impedes DE establishment (Nord et al., 2020). We therefore hypothesized that a second fate choice between mesodermal and endodermal fates further exacerbated the uneven mesodermal fate preference between the treatments. Indeed, our analysis indicates that in the mesodermal-endodermal fate choice, the majority of WAG treated cells differentiate toward mesodermal fate. This is in contrast to the WAN and WA treatment in which cells preferentially differentiate toward definitive endoderm. WAB treated cells exhibited mixed potential, and split roughly equally along the two paths.

We sought to elucidate a possible explanation for Foxo1’s role in this fate choice by searching for the shortest paths toward a set of endodermal genes. Specifically, similar to the previous mesodermal analysis, we applied community detection to the full endodermal network and identified the cluster containing Sox17 and Foxa2, known master regulators of endodermal fate (**Figure 7c**). This cluster included the transcription factors Sox17, Foxa2, Bmp2, Cited1, Foxa1, Gata6, Hhex, Lhx1, and Tcf7l2. For each treatment-specific network, we computed the shortest paths from Foxo1 targets (as determined in the same manner as the previous Wnt analysis) to these genes (**Figure 7d**). Importantly, we found that many of these regulators, including both Sox17 and Foxa2, were not reachable from Foxo1 in the WAG network, consistent with the observation that WAG treated cells preferentially differentiated toward mesodermal fate over endodermal fate. Interestingly, Foxa2 was not reachable in the WAN network, though Sox17 was. We believe this may explain the differences in distribution of cells between the endodermal clusters 7,1 and 7,0. Namely, of the cells that specify endodermal fate, WA and WAB treated cells more fully transitioned into cluster 7,0. In contrast, WAN treated cells remained mostly in cluster 7,1 and were less likely to differentiate further into 7,0. Previous studies in directed differentiation of human ESCs toward DE have implicated Sox17’s regulatory role in establishing DE to be upstream of Foxa2, the loss of which impairs foregut and subsequent hepatic endoderm differentiation (Genga et al., 2019). This is consistent with our cell type annotations of cluster 7,1 (definitive endoderm) and 7,0 (gut endoderm), and offers a possible mechanism of the discrepancy in cell fate preferences between treatments. Taken together, these results demonstrate that WAB, WAN, and WA networks do not allow for activation of mesoderm programs from their cognate effectors, but instead assume topologies that are preferential for endodermal fate, implying profound structural changes in the networks that underlie the treatment-specific differences we see in cell fate. Further, this analysis provides a basis for identifying which signaling pathways must be targeted for directing certain fate transitions, for example through an exhaustive search of effector targets.

### Target gene differences amongst TFs confirm network topology restructuring dependent on treatment

Given the apparent restructuring of GRN topology upon modulation of signaling activity, we aimed to understand the extent to which broad, network-wide topology changes were responsible for differences in cell fate potential between treatments. To answer this, we performed two distinct analyses focusing on differences in targets of transcription factors across the four treatment networks.

First, we asked if transcription factors actively expressed in all four treatments regulated the same set of target genes. To this end, we focused on the treatment-specific networks along the mesodermal path. We narrowed our analysis to the 72 TFs that were active in all four treatments during day 3 or day 4 along the mesodermal path. We quantified overlap between targets of each TF in a pairwise comparison of the treatments using the Jaccard similarity (**Figure 8a**). As a baseline, we implemented a bootstrapping method in which we reconstructed 10 networks for each treatment along the mesodermal path (using 400 sampled cells each); for each TF we calculated the average Jaccard similarity of its targets amongst pairwise comparisons of the reconstructed networks in a treatment. This gave us a baseline, or expected, Jaccard similarity for each TF in each treatment. Overall, we found that overwhelmingly, target differences between treatments were significantly greater than the baselines, indicating vast network topology differences amongst the treatment networks.

**Figure 8.**
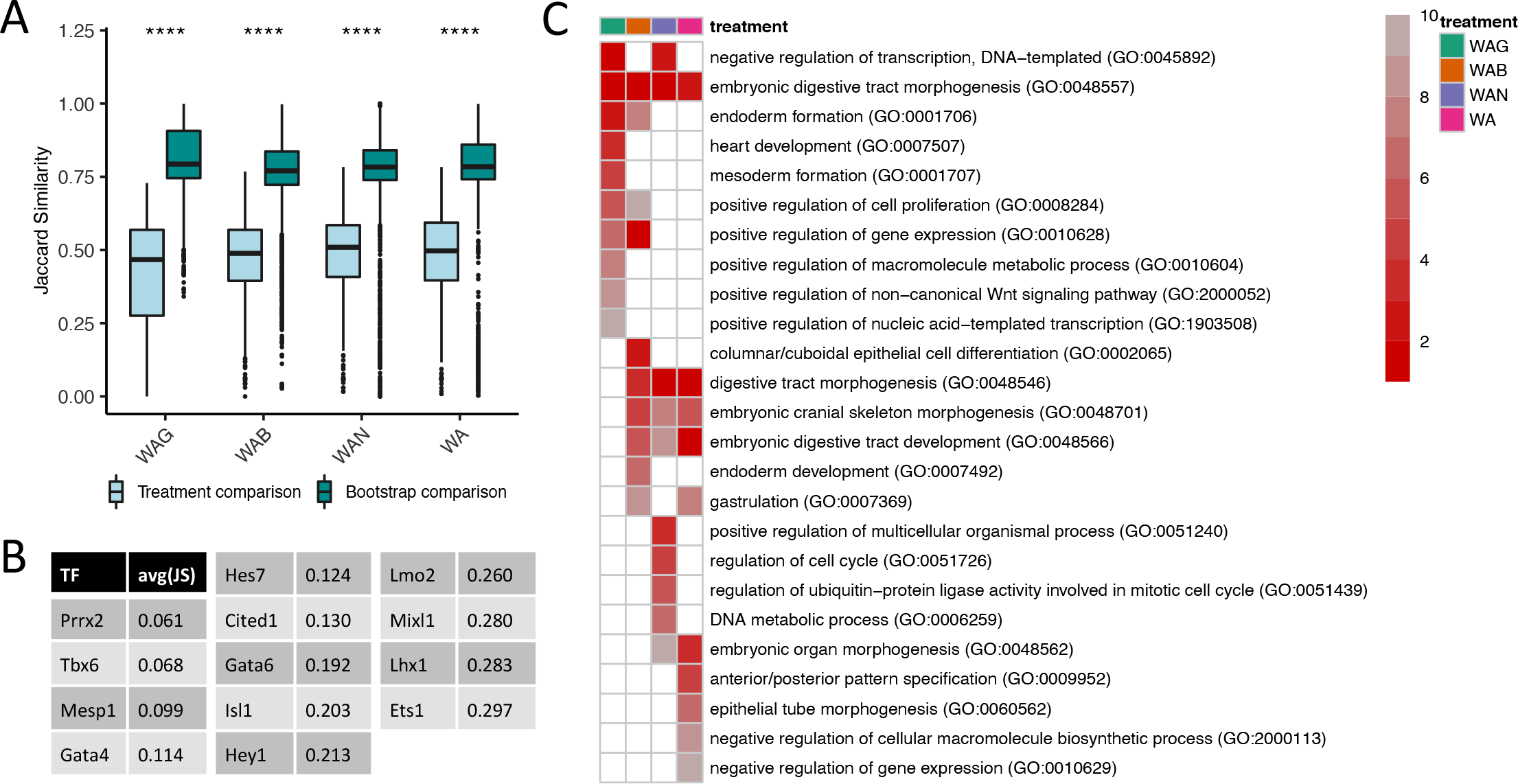
Similarity of targets of TFs in different treatments. (A) Jaccard similarity of targets of 72 TFs. The targets of each TF in a treatment are compared to those in the other three treatments (light blue). As a baseline (turquoise), targets of each TF in bootstrapped reconstructed networks within a treatment are pairwise compared. All pairs p<2×10^-16^. (B) The 13 TFs selected for GSEA and their average pairwise Jaccard similarity amongst the treatment-specific networks. (C) Summarized results of performing GSEA on targets of TFs in B. Heatmap shows rankings (top 10 shown) based on frequency a term is considered enriched amongst the 13 TFs (1 = most frequent enriched term).

Next, we asked if these differences in predicted targets had functional consequences in impacting fate potential. To this end, we focused on the 13 TFs amongst the 72 mentioned above that exhibited the largest differences in predicted targets between treatments (**Figure 8b**). For each treatment, we performed gene set enrichment analysis on the predicted targets of these TFs (**Figure 8c**). Interestingly, we found enrichment for heart development and mesoderm formation in only the WAG treatment, despite the fact that the networks we analyzed were reconstructed along the mesodermal path. Of note, all treatments showed enrichment for embryonic digestive tract morphogenesis. Importantly, prior literature has indeed hinted at environment-based differences in fate potential. For example, in uncovering the regulatory structure underpinning naïve pluripotency in mESCs, the effect of Esrrb/Nanog double knockdown was shown to be dependent solely on culture environment with self-renewal impeded in the presence of 2i alone, but not in the presence of both 2i and LIF (Dunn et al., 2014). Taken together, these results support our hypothesis that manipulating signaling activity results in a topological restructuring of the GRN, ultimately guiding cell fate potential.

### Patterning programs and residual neuroectoderm programs drive differences between in vivo and in vitro gastrulation and mesoderm specification

Finally, we aimed to understand the extent to which our *in vitro* mESC-derived mesodermal cells established a GRN resembling that of their *in vivo* counterparts. To this end, we turned to a mouse gastrulation dataset from Grosswendt et al., and sampled 250 cells from each of four annotated populations that corresponded to our *in vitro* mesoderm lineage populations: “Epiblast”, “Primitive streak early”, “Primitive streak late”, and “Mesoderm presomitic”. We then used Epoch to reconstruct an *in vivo* dynamic network from this sampled data. After reconstruction, we extracted the top regulators in each epoch by both PageRank and Betweenness-Degree (**Figure 9a**). Interestingly, there was substantial difference between the top regulators of the *in vivo* and *in vitro* networks, suggesting broader topology differences between the two. Despite this, some key regulators were shared, including Peg3, which ranked highly in the second epoch of the *in vivo* network in both PageRank and Betweenness-Degree. This further supported our earlier results implicating a role for Peg3 in orchestrating the pluripotent to mesodermal fate transition *in vitro* and *in vivo*.

**Figure 9.**
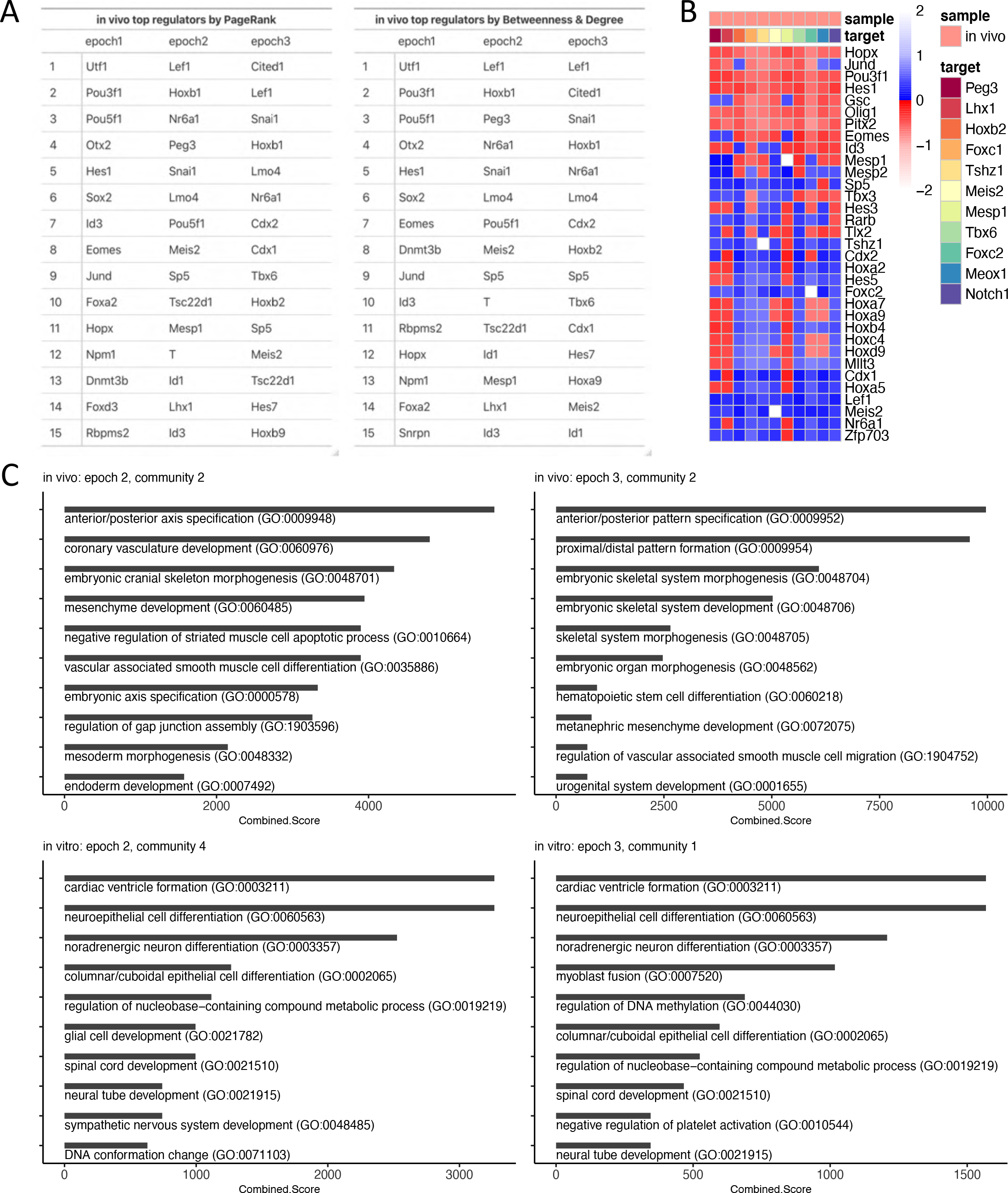
Comparison to *in vivo* gastrulation and mesoderm specification. (A) Top regulators of each epoch in the *in vivo* mesoderm network as predicted by PageRank and Betweenness-Degree. (B) Shortest path analysis showing length of shortest paths from Wnt effector targets (each row) to mesodermal target genes (each column) in the *in vivo* network. Path lengths are normalized against average path length in the network. Blue and red indicate paths that activate and repress the target gene respectively. If such a path does not exist, it has length of infinity and is white. (C) Top ten enriched terms for *in vivo-* and *in vitro*-specific modules based on the Combined Score from Enrichr GSEA analysis.

Of the 22 unique TFs in the second and third epochs of our top regulator lists, at least 17 have known roles in guiding exit from pluripotency, mesoderm specification, or somitogenesis. In particular, a number of top regulators in the second epoch are required for guiding the exit of pluripotency, such as Nr6a1 and Snai1 (Galvagni et al., 2015; Gu et al., 2005). Others in this epoch are components of the Wnt signaling pathway including Lef1, a Wnt effector, and Sp5, which itself is directly activated by Wnt signals and further aids in activating Wnt target genes to promote differentiation into multipotent mesodermal progenitors (Kennedy et al., 2016). Further, TFs like brachyury and Lhx1 are necessary for primitive streak formation and normal development of primitive streak-derived tissues (Shawlot et al., 1999; Wilson et al., 1995). Spread across both the second and third epochs are a number TFs responsible for patterning and promoting mesoderm fate. These include TFs like Hoxb1, Hoxb2, Hoxb9, Cdx1, Cdx2, Mesp1, Meis2, and Id1 (Bernardo et al., 2011; Cecconi et al., 1997; Foley et al., 2019; Gouveia et al., 2015; Malaguti et al., 2013; Roux et al., 2015; Takahashi et al., 2005, 2007). Of these, many are also responsible for inhibiting alternative fates, such as Id1’s role in inhibiting neural induction and Cdx2’s role in blocking cardiac differentiation (Mendjan et al., 2014; Zhang et al., 2010). Finally, at least two TFs in the third epoch, Tbx6 and Hes7, are essential for specifying and coordinating somitogenesis (Bessho et al., 2001; Chapman et al., 2003). Ultimately, the predicted top regulators match well with prior literature describing the ESC to presomitic mesoderm lineage trajectory from which the network was reconstructed. Additionally, this further corroborates Epoch’s usefulness and ability in isolating important genes driving dynamic processes.

We further applied the signaling cascade tracing analysis to the *in vivo* network. Analogous to the *in vitro* MULTI-seq network, we looked for paths connecting Wnt effectors to the same TFs in the *in vitro* WAG-specific modules (**Figure 9b**). As expected, all of the TFs were reachable. Further, Wnt activation and these target TFs were highly connected, with paths existing from almost every effector to every TF, implying the existence of tight and coordinated control over this mesoderm module.

Finally, we sought to directly compare the topological and functional differences between the *in vivo* and *in vitro* mesoderm networks. Because the data originates from separate experiments, to directly compare topological differences we applied a threshold to both networks, keeping the top 2% of non-zero-weighted edges in each. We then extracted the differential network corresponding to *in vivo*-specific interactions and the differential network corresponding to *in vitro*-specific interactions (**Supplementary Figure 12**). We performed community detection on both differential networks and measured the activity of each resulting module by assessing the average expression of member genes across time in the *in vivo* and *in vitro* data (**Supplementary Figure 13a,b**).

Of the *in vivo*-specific modules, three were not activated or were insignificantly activated in the *in vitro* data, despite being strongly activated in the *in vivo* data. To understand the functional consequences of this, we applied GSEA to each of these modules (**Figure 9c, Supplementary Figure 13c**). Two of these, one from the second epoch and one from the third epoch, showed enrichment for multiple pathways related to patterning and axis specification, including anterior/posterior axis specification, anterior/posterior pattern specification, proximal/distal pattern formation, and embryonic axis specification, amongst others. Thus, a large, but not unexpected, difference between the *in vivo* and *in vitro* mesoderm is the lack of activation of patterning programs in the *in vitro* ESC-derived mesodermal cells.

Conversely, of the *in vitro*-specific modules, we isolated those that were not activated or were insignificantly activated in the *in vivo* data but strongly activated in the *in vitro* data. Within the first epoch, three communities satisfied this criterion. We hypothesized that differences in this early epoch could drive fate differences between the two networks at later time. GSEA on these modules revealed that one was enriched for positive regulation of stem cell proliferation and regulation of mRNA splicing while another was enriched for terms related to fluid shear stress, DNA methylation, and male gonad development (**Supplementary Figure 14**). Meanwhile, GSEA on in vitro-specific modules from the second and third epochs revealed the underlying activation of neural-related programs in the *in vitro* ESC-derived mesoderm (**Figure 9c, Supplementary Figure 13d**). Most of the modules were enriched for neuroectoderm processes such as neuroepithelial cell differentiation, noradrenergic neuron differentiation, glial cell development, spinal cord development, and forebrain neuron differentiation, amongst others. Importantly, this implies that the *in vitro* differentiated cells failed to completely inhibit the default neuroectoderm lineage and instead retained a network topology capable of activating at least portions of neuroectoderm programs. This aligns with our observation that the vast majority of the *in vitro* differentiated cells tended toward the neuroectoderm lineage, with only a small percentage escaping this fate and specifying for mesoderm. Taken together, our results suggest that to produce more faithful mesodermal cells more efficiently, emphasis should be placed preventing or disrupting network module topologies responsible for neural programs.

## Discussion

Methods of inferring regulatory network structures are essential for understanding cell identity transitions, uncovering mechanisms of disease progression due to dysfunctional gene regulation, and answering other regulatory questions in biology. Unfortunately, experimental methods are unscalable and existing computational methods suffer from poor performance or high computational burden. Additionally, these approaches typically infer static networks, which do not elucidate the network topology transitions that drive dynamic processes such as differentiation. Prior work addressing this has involved utilizing a combination of bulk RNA-Seq, TF ChIP-Seq, and Pol2-ChIA-PET data, with the aim of identifying differences in GRN activity or structure at discrete experimental time points during the shift from naïve to formative pluripotency (Kim et al., 2020). However, such an approach is limited in resolution and scale due to the discrete nature of the data and the sparsity of cell type-specific ChIP-Seq data, ultimately resulting in networks centered around just a handful of TFs. Thus, an approach that relies solely on single-cell transcriptomics is an attractive solution, as single-cell trajectory inference tools are capable of organizing individual cells along a continuum, improving resolution and accounting for asynchronous processes. For those reasons, we developed a novel GRN reconstruction tool, Epoch, which leverages single-cell transcriptomics and efficiently infers dynamic network structures. We demonstrate that Epoch outperforms CLR, GENIE3, and existing single-cell GRN reconstruction tools in both synthetically generated and real-world datasets. Furthermore, Epoch’s relative efficiency allows it to be easily applied iteratively for applications requiring fine-tuning of network topology or iterative simulations of changes in network structure.

Epoch’s utility is additionally enhanced by its flexibility. There are no strict requirements on pseudotemporal input as we demonstrate here using synthetic simulation time, diffusion pseudotime, and RNA velocity-based latent time. We further validate on the *in vivo* muscle development dataset that precision-recall is largely comparable for different pseudotime methods (**Supplementary Figure 1f,g**). Additionally, Epoch’s workflow is structured in discrete steps allowing users to pick-and-choose as well as substitute portions of the workflow. We anticipate that this flexibility will also be useful as advances in sequencing technology, such as multi-modal methods, and analysis techniques, such as novel cell ordering or reconstruction methods, can be easily adapted into Epoch’s framework. Importantly, because of single-cell transcriptomic data’s inherent noisiness, we also asked if smoothing gene expression over time could alleviate noise-related drops in reconstruction performance. We compared performance between unsmoothed and kernel-smoothed data and found that the there was no significant difference (**Supplementary Figure 1h**). However, smoothing functions remain in Epoch’s code to aid in cleaner visualizations of gene dynamics.

Finally, we applied Epoch to scRNA-seq data from day 0 through day 4 *in vitro* mESC directed differentiation to uncover underlying dynamic networks driving mesodermal lineage specification. Our data consisted of cells transitioning into all three germ layers, and Epoch was able to recover a number of key lineage drivers. Because our data consists of cells undergoing four different treatments (WAG, WAB, WAN, WA), we were additionally able to reconstruct treatment-specific GRNs. Our analysis uncovered significant treatment-induced topological restructuring that altered the lineage specification landscape, resulting in treatment-based differences in neuroectoderm, mesoderm, and endoderm reachability. Specifically, in direct contrast to the cells in the WAB, WAN, and WA treatment groups, the cells in the WAG treatment group had higher mesodermal fate potential, exhibited a relatively stunted neuroectodermal trajectory, and had decreased potential for endodermal fate. Importantly, we uncovered three sets of topological differences that may play a role in specifying mesodermal fate.

First, we identified a set of TF modules that were preferentially activated in WAG treated cells compared to cells undergoing the other three treatments along the mesodermal fate. Specifically, these modules encompassed the transcription factors Peg3, Lhx1, Hoxb2, Foxc1, Tshz1, Meis2, Mesp1, Tbx6, Foxc2, Prrx2, Meox1, and Notch1. Upon further analysis, we observed that the WAG network developed a topology that promoted Peg3 activity, which was predicted by Epoch to have a role in regulating a large fraction of these mesodermal TFs. This influence was again specific to WAG treated-cells, and indeed, prior studies have implicated a GSK3-dependent pathway for the epigenetic regulation of imprinted loci including Peg3 (Meredith et al., 2015). Peg3 expression has been detected in early somites and other mesodermal tissues (Kuroiwa et al., 1996). Interestingly, it has been separately demonstrated that knockdown of Peg3 in mouse embryonic fibroblasts and neural stem cells increases expression of pluripotency genes and enhances overall reprogramming efficiency (Theka et al., 2017). Taken together these results suggest a role for Peg3 in modulating the exit of pluripotency and mesodermal lineage specification. If so, this finding could paint Peg3 as a highly enticing transcription factor to target with the goal of improving efficiency of directed differentiation protocols toward certain mesodermal fates such as somites.

Next, we compared topological differences in the treatment-specific networks by examining the existence of, and computing the length of, shortest paths. Specifically, we sought to trace paths from signaling effectors to the activation of mesodermal genes within these dynamic networks, thereby providing possible mechanisms of how signal transduction pathways not only directly regulate targets corresponding to distinct genetic programs, but also restructure network topology to alter accessibility to targets, which hones the capacity and potential for distinct fates. A number of studies have pointed to the influence of “Wntch” signaling on the neuroectoderm-mesendoderm cell fate decision in gastrulation (Hayward et al., 2008). Our data was consistent with these reports: WAB, WAN, WA treated cells exhibited activated Notch pathways and exhibited full trajectories toward neuroectoderm fate; WAG treated cells exhibited markedly diminished or non-existent Notch signaling and reached mesodermal fates while seemingly displaying a stunted capacity for neuroectodermal fate. Beyond this, we demonstrated that WAG treated cells established a GRN topology that allowed for Wnt-dependent mesoderm specification. Specifically, our analysis indicated the existence of paths from Wnt signaling effectors to the aforementioned mesodermal TFs within the WAG network. Analogous paths were markedly sparser in WAB, WAN, and WA networks, and some mesodermal TFs were entirely unreachable from Wnt effectors. This further supported our hypothesis that signaling-induced topological differences altered the cell fate landscape, resulting in distinct propensities for specific fates, though the nature of the emergence of these topological differences remains unclear. One possibility is that the network structures are dynamically altered by epigenetic events initiated through the various signaling pathways activated in treating the cells. We point to GSK3’s reported effect over Peg3 methylation as an example of such a phenomenon. Further studies to elucidate differences in chromatin accessibility between mESCs undergoing directed differentiation by way of distinct treatments, or activation of distinct signaling pathways, will aid in clarifying if such a mechanism is at play.

We then focused our analysis on cells specifying mesendoderm fate, and outlined topological differences that could lead to differences in mesoderm vs. endoderm fate potential in cells undergoing different treatments. We applied an analogous shortest path analysis along the endodermal fated path to identify the link between suppression of PI3K signaling, subsequent Foxo1 activation, and endoderm specification. We again found topological differences between WAG and the remaining three treatments, with no paths from Foxo1 to essential endodermal regulators such as Sox17 and Foxa2 in the WAG network, explaining the preference of these cells to transition into mesodermal rather than endodermal fates. Interestingly, we found that Foxa2 was not reachable in the WAN network, which was consistent with our finding that of the WAN treated cells that specified endodermal fate, the majority tended to remain in earlier endodermal stages based on RNA velocity. This was in contrast to WAB and WA treated cells, which tended to transition more fully into a later endodermal stage, thereby explaining the observed differences in mesodermal vs. endodermal fate potential between the treatments.

Finally, we compared the network driving *in vitro* directed differentiation toward mesoderm fate against the network driving the analogous *in vivo* cell identity transition. Direct comparison of the two networks based on predicted top regulators and the extraction of edge-level differential networks suggested broad differences in the topologies of the established networks. Importantly, community detection and GSEA revealed two major differences. First, the *in vivo* network established modules related to patterning and axis specification programs that were not established in the *in vitro* network. This is unsurprising given that while embryoid bodies exhibit basic self-organization, they do not fully recapitulate the self-organization present in *in vivo* gastrulation (ten Berge et al., 2008; Glykofrydis et al., 2021). It is further possible that failing to establish these patterning modules hindered the *in vitro* cells from establishing network topologies conducive for mesoderm fate (or specific downstream mesodermal fates), limiting their differentiation potential. Second, the *in vitro* network retained a topology at least partially favorable for neuroectoderm development, as there were multiple modules enriched for neural-related processes in the *in vitro* network. This suggests that the cells undergoing *in vitro* directed differentiation incompletely suppressed the neuroectoderm fate, and further corroborates our observation and provides explanation for why the *in vitro* differentiating cells overwhelmingly specified for the neuroectoderm path. Overall, this analysis suggests that guiding cells toward a more faithful mesodermal fate at a higher rate will likely require steps to disrupt formation or retention of neuroectoderm-favoring network topologies while possibly promoting the establishment of patterning-related programs.

Our results ultimately suggest a model of differentiation that is driven by activation of genetic programs, but honed by network topology changes. In other words, it is likely that the signaling-induced restructuring of the GRNs alters the fate potential landscape which allows for independent control of multiple genetic programs, increasing the diversity of reachable cell states. In the case of our mESC directed differentiation, GRN topology clearly restricted the potential of cells toward specific fates, resulting in different propensities for mesoderm, endoderm, and neuroectoderm depending on treatment. Finally, while it may be the case that both signaling activities described (i.e. the effector regulation of distinct targets and the restructuring of network topology) occur concurrently, it is ultimately difficult to resolve their relative timing.

We believe the single-cell directed differentiation dataset collected in this study will be a useful resource for further analysis of decision-making in early gastrulation. Furthermore, the methodologies presented here, available in the R package Epoch, are broadly applicable to any biological question that may benefit from uncovering dynamic regulatory network structures and the comparison of such networks. In particular, we can apply Epoch to understand cell fate transitions and decisions at branch points in lineage trajectories, uncover key regulators driving these decisions, and identify paths from the signal transduction level to the transcriptional regulation level. Such an approach provides a powerful strategy to not only elucidate dynamic, multi-scale processes in development, but also to identify signaling pathways to dysregulate for the purposes of directing cell identity transitions *in vitro*.

## Acknowledgements

We would like to thank Yuqi Tan, Ray Cheng, Eric Kernfeld, and David Johanson for helpful discussions and testing Epoch code. We also thank Chris McGinnis and the Gartner Lab for their help implementing the MULTI-seq protocol and for providing the required reagents. This work was supported by the National Institutes of Health under grant R35GM124725 to PC and by the National Science Foundation Graduate Research Fellowship under Grant No. DGE-1746891 to EYS.

## Methods

### mESC maintenance and differentiation

GFP-Brychyury mESC cells (Gadue et al., 2006) were maintained and differentiated through day 2 of the directed differentiation protocol according to Spangler et al. (Spangler et al., 2018). At day 2, four different primitive-streak induction treatments were established by adding growth factors Wnt3a (25ng/mL) and Activin A (WA; 9ng/mL) along with one of the following additional growth factors: Noggin (150ng/mL), Gsk inhibitor (10mM), or BMP4 (0.5ng/mL). On the day of sequencing, EBs and monolayers were dissociated to a single cell suspension through the use of TrypLE and 40uM cell strainers.

### Multi-Seq protocol

We utilized the MULTI-seq sample barcoding and library preparation protocol from the McGinnis lab (McGinnis et al., 2019) in order to sequence 27 samples in two 10x capture runs. In short, the protocol involved tagging cell membranes with sample specific barcodes using a lipid-modified oligonucleotide (LMO). The LMOs (reagents obtained from the McGinnis lab) anchor into the cell membrane and allow for the attachment of a normal ssDNA oligonucleotide (MULTI-Seq barcode). A unique MULTI-seq barcode was added to each sample after which, all samples were pooled and sequenced as if they were one sample. An equal number of cells from each sample were combined to make up the pooled samples and ensure equal representation of each sample after down-stream sequencing. MULTI-Seq barcodes were used down-stream to identify which cells were from which samples.

### Library preparation and sequencing

After cells were successfully tagged and pooled, they were submitted to the sequencing core facility for 10x capture and library preparation. A Truseq library preparation was performed with the necessary adjustments made to accommodate the MULTI-Seq platform. Libraries were sequenced on Illumina NovaSeq.

### Single-cell data processing

Sequencing alignment to the mm10 reference genome was performed using CellRanger (version 3.1), and spliced, unspliced, and ambiguous read counts were extracted using Velocyto. Sample barcode classification was performed using the deMULTIplex R package (https://github.com/chris-mcginnis-ucsf/MULTI-seq). Subsequent quality control filtering, normalization, clustering, and differential gene expression analysis was performed using Scanpy (version 1.4.5) (Wolf et al., 2018). Specifically, doublets and negatives from the sample barcode classification were removed, along with 271 cells we identified as residual fibroblasts (Thy1+). Next, genes were excluded if they were detected in less than 5 cells; cells were excluded if their mitochondrial gene content exceeded 10% of their total reads or if they had fewer than 500 unique genes. This left us with 5530 cells and 17528 genes. The data was then normalized and log transformed before highly variable genes were identified (1938 genes). The data was scaled and PCA was performed. Leiden clustering was performed and visualized on a UMAP embedding. We further removed a cluster of possible dying cells expressing relatively high mitochondrial gene content and low total read count, leaving a dataset with 5228 cells and 1938 variable genes. Leiden clustering was refined, and RNA velocity was computed using scvelo (version 0.2.2) (Bergen et al., 2020). Latent time was computed with scvelo, using root and end states that were computed through CellRank (version 1.0.0-rc.0). Cell type classification was performed via SingleCellNet (Tan and Cahan, 2019), using complied embryo data for training the classifier. For the E12.5 muscle development data and day 4 early directed differentiation data, diffusion maps and diffusion pseudotime were computed through Scanpy.

### Synthetic benchmarking

15 synthetic datasets were generated using the Dyngen package in R (Cannoodt et al., 2020). Specifically networks were generated with 100-420 genes (including 20-70 TFs), containing varied topological motifs, and cell trajectories were simulated over time. Simulated single-cell sequencing was performed, resulting in datasets that included up to 1000 cells. CLR was implemented with the minet package in R (Meyer et al., 2008), and GENIE3 was implemented with the GENIE3 package in R (Aibar et al., 2017; Huynh-Thu et al., 2010). Variations of these two methods were carried out using the Epoch framework. Performance (AUPR) was compared against sets of 5 random networks generated for each reconstructed network via the random permutation of edge weights across targets.

### BEELINE benchmarking

The BEELINE benchmarking tool (Pratapa et al., 2020) was used to compare Epoch against the following single-cell reconstruction methods (using the pre-built Docker images from BEELINE): GENIE3, GRISLI, GRNBoost2, GRNVBEM, LEAP, PIDC, PPCOR, SCODE, SCRIBE, SINCERITIES, and SINGE. We added the four versions of Epoch into the BEELINE framework for benchmarking. The synthetic and curated datasets (including simulation time and pseudotime) on which we did the benchmarking were the same as those used in the BEELINE publication, and downloaded from https://doi.org/10.5281/zenodo.3378975. Briefly, for synthetic datasets, each trajectory type included 5 dataset sizes (100 cells, 200 cells, 500 cells, and 2000 cells), and each dataset size included 10 distinct datasets (for a total of 50 datasets for each trajectory type). Curated datasets were simulated from four literature derived networks (GSD: gonadal sex determination, HSC: hematopoietic stem cell differentiation, VSC: ventral spinal cord, mCAD: mammalian cortical area development). 10 datasets were simulated for each network at 3 different dropout levels (0%, 50%, 70%), for a total of 30 datasets per network. AUPR, early precision, and runtimes were evaluated. BoolODE (Pratapa et al., 2020) was used to re-simulate the curated datasets for the purposes of extracting the true simulation time.

### Bootstrapped network reconstruction of early MULTI-seq data

We isolated early latent time clusters 1 and 3 and uniformly sampled 400 cells, 10 times to reconstruct 10 networks. For each reconstructed network, we extracted top regulators by PageRank and identified TFs that were most frequently ranked in the top 10 regulators. We further computed the average normalized betweenness and average normalized degree across the 10 reconstructed networks to identify top regulators.

### Differential network analysis and assessment

As one method of network comparison, we used a naïve edge weight-based approach to extract the most differentially weighted edges between networks. Briefly, this involves computing edge weight differences for each possible edge in the two networks. Edges are then considered “differential” and specific to a given network if the edge weight difference is greater than a given threshold (in the case of both the treatment difference analysis and the in vivo vs. in vitro comparison, this threshold was set to 2). Such an approach resulted in “differential networks” which are representative of unique topological features existing in each treatment-specific network. To assess the feasibility of this simple method in extracting real and important topological differences between networks, we first applied the same method to compare the three lineage networks in the dataset. The ground truth on which we assessed these differential networks was a set of lineage-unique “pseudo-ChIP-seq” networks we compiled similar to TF-promoter binding predictions generated by (Lu and Mar, 2020) (**Supplementary Figure 6a**). In short, TF-target interactions were acquired by mapping the PWM of a TF (Weirauch et al., 2014) to the promoter region sequence of potential target genes using the Find Individual Motif Occurrences (FIMO) software (described in more detail below). These ground truth networks were filtered for dynamically expressed genes along given paths (for lineage-specificity), and further filtered for unique edges to mimic extraction of unique topological features (lineage-unique). Using this set of networks, we compared AUPR of our path-differential networks against random networks (generated by shuffling edge weights of targets of each TF in the differential networks). On average the differential networks had a 1.7 fold improvement over random AUPR, with many comparisons falling in the 1.8-3.0 range (**Supplementary Figure 6b**).

### Construction of path-unique TF-target pseudo-gold standard

Transcription star site (TSS) annotations were downloaded from the USCS Genome Browser (mouse GRCm38/mm10 genome assembly) (Haeussler et al., 2019). Promoter regions were defined as -750 to +250 around the TSSs. We then acquired position weight matrices (PWMs) of transcription factors from the Catalog of Inferred Sequence Binding Preferences (CIS-BP Database build version 2.00) (Weirauch et al., 2014). We used the Find Individual Motif Occurrences tool (Grant et al., 2011) to obtain potential TF-promoter binding pairs for 648 TFs. We kept all interactions for which p-value < 5 ×10^-5^. To generate path-unique gold standards, for each lineage (mesoderm, endoderm, neuroectoderm) we filtered interactions for dynamically expressed genes along each path as computed by Epoch. Further, when assessing the “differential network” pairwise comparisons, we filtered out interactions that were redundant between the gold standards, and only kept interactions unique to each path.

### Targets of TFs by treatment comparison

To quantify the extent of topology differences between treatments along the mesodermal path, we performed pairwise comparison on the four treatment-specific networks and computed Jaccard similarity between predicted targets of TFs. The comparison was carried out on 72 TFs that were selected based on their active expression on day 3 or day 4 (i.e. after induction treatments). We computed baseline similarities for each TF in each treatment by bootstrapping the network reconstruction (10 network reconstructions per treatment, 400 sampled cells per network reconstruction) and averaging their pairwise Jaccard similarities. We selected the 13 TFs exhibiting the greatest differences in targets amongst the treatments and performed gene set enrichment analysis using the Enrichr R package (Chen et al., 2013; Kuleshov et al., 2016) and the GO Consortium Biological Processes database (Ashburner et al., 2000; Gene Ontology Consortium, 2021) on the targets of each TF. We further filtered out terms that did not meet the criteria of adjusted p-value<0.05. Finally, we summarized the results between the 13 TFs by counting the frequency (out of 13) that a term was considered enriched and ranked the results accordingly.

### Comprehensive signaling effector TF targets list

We acquired binding score (MACS2) data for 18 signaling effector TFs from the ChIP-Atlas (Oki et al., 2018), and processed each accordingly: target genes were ranked by maximum binding score and the top 2000 targets were retained (or all retained, if less than 2000 targets). To compute average effector activity, target lists were filtered for those appearing in the MULTI-seq directed differentiation data, and not considered in the analysis if the effector had less than 10 targets after filtering.

### In Vivo Comparison

To create a comparable in vivo dataset, 250 cells were randomly sampled from each of the following annotated populations in the Grosswendt et al. gastrulation dataset: “Primitive streak anterior”, “Endoderm”, “Ectoderm early 1”, “Ectoderm early 2”, “Epiblast”, “Fore_midbrain”, “Future spinal cord”, “Endoderm gut”, “Neural crest”, “Ectoderm neural anterior”, “Ecotderm neural posterior”, “Neuromesodermal progenitor late”, “Node”, “Notochord”, “Mesoderm pharyngeal arch”, “Mesoderm presomitic”, “Primitive streak late”, “Primitive streak early”, “Somites”, “Neuromesodermal progenitor early.” The resulting dataset spanned E6.5 to E8.5. Quality control filtering, normalization, and Diffusion Pseudotime trajectory inference was performed using Scanpy (version 1.4.5) (Wolf et al., 2018), analogous to the processing of the MULTI-Seq data described above. Further comparison was limited to the mesodermal lineage – this included populations labeled “Epiblast”, “Primitive streak early”, “Primitive streak late”, and “Mesoderm presomitic”.

Epoch was employed to reconstruct the in vivo dynamic network corresponding to this data. To compare the *in vivo* network to the *in vitro* network, both networks were thresholded such that the top 2% of non-zero edges were kept. Network comparison including differential network extraction and community detection were performed using Epoch. Average module expression was analyzed by computing mean expression of predicted activated members over time if the module included at least 8 such members (members predicted to be repressed were not included in the average so as not to improperly depress this measure of activity). GSEA was performed using the Enrichr R package (Chen et al., 2013; Kuleshov et al., 2016) and the GO Consortium Biological Processes database (Ashburner et al., 2000; Gene Ontology Consortium, 2021). Both the 2021 and 2018 versions of this database were used for enrichment analyses. Figures shown here show results using the 2021 version of this database for all enrichment analyses except for epoch3, community 3. This community was not enriched for any terms in the 2021 database, but was enriched for multiple terms in the 2018 pathway.

### Epoch Workflow

Epoch takes as input processed (normalized and log-transformed) single-cell transcriptomic data and accompanying pseudotime (or equivalent) annotation. The Epoch workflow is based in three strategies: (1) the extraction of dynamically expressed genes and subsequent static reconstruction with CLR, (2) network refinement using a cross-correlation based strategy termed “cross-weighting,” (3) the extraction of a dynamic network and subsequent identification of top regulators. Epoch is available as a package in R, and code and tutorials can be found at https://github.com/pcahan1/epoch.

### Identification of dynamically expressed genes

Limiting network reconstruction to dynamically expressed genes serves two purposes. Importantly, it focuses the network on interactions that are more likely playing a role in the observed biological process. Second, it reduces the instances of false positive interactions and improves precision by limiting possible edges between temporally-variable genes. To select for such genes, Epoch models individual genes across the annotated pseudotime using a generalized additive model (GAM). Specifically, Epoch uses the ‘gam’ package (gaussian family, loess smooth pseudotime) to fit a model for each gene using the backfitting algorithm. Genes are considered dynamic based on significance of the smooth term (pseudotime). Alternatively, users can specify to find dynamic genes via TradeSeq (Van den Berge et al., 2020), a recently developed tool that identifies dynamic changes in gene expression via a GAM based on the negative binomial distribution.

### CLR

The Context Likelihood of Relatedness (CLR) algorithm was first proposed in 2007 by Faith et al. as a method for inferring gene regulatory networks from expression data. In short, CLR computes the mutual information (MI) between expression levels of every potential transcription factor and target gene. From here the method scores the MI values in the context of the network by computing a z-score relying on (1) the distribution of MI values for all potential regulators of the target gene, z_i_, and on (2) the distribution of MI values for all potential targets of the transcription factor,

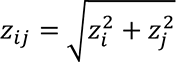

The result is a weighted adjacency matrix describing the likelihood of TF-target gene interactions that can be thresholded to derive a network structure. For Epoch, a version of CLR is applied to dynamically expressed genes to reconstruct an initial static network via the ‘minet’ package in R. Users can choose between implementing the MI version or a version using Pearson correlation in place of MI. In either case, Epoch at this step will return a GRN table.

Each row outlines an interaction in the initial network identifying the transcription factor, target gene, correlation, and CLR-based z-score.

### Cross-weighting

After inferring an initial network structure, Epoch can apply a network refinement step we call cross-weighting. The objective of this step is to negatively weight edges in the initial network that are unlikely to be true interactions, that may, for example, be representative of indirect interactions. To this end, for every TF-target pair in the initial network, Epoch computes cross-correlation across a given lag time (defaults to a fifth of total pseudotime). After ordering the lag times by decreasing correlation, Epoch computes an offset value where maximum correlation is achieved, defined by default as the top third of the ranked lag times. Finally, Epoch scores the offset values of each interaction:

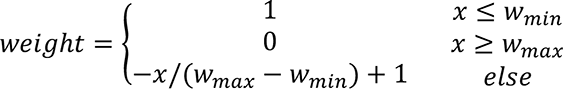

where the offset, x, is computed as described, and maximum and minimum windows can be altered by the user if desired. Z-scores from the initial network are weighted accordingly, ultimately filtering out false positives. Epoch, at this step will return an appended GRN table, including for each interaction, the offset value and new weighted score.

Default parameters for cross-weighting are specified within Epoch, and were chosen based on empirical improvements in performance across synthetic and real datasets (ranging in number of genes, number of cells, and simple trajectory types). For example, we found that optimal lag time varied to some extent with dataset size, with larger datasets requiring a larger lag time to catch target response to regulator expression changes (**Supplementary Figure 15a**). Smaller lag times tended to correlate with decreased AUPR, likely because the lag time was not sufficient to catch the point of maximum cross-correlation. AUPR also begins to gradually decrease at larger lag times, though this effect is much less pronounced. We found that our default lag (set to 1/5 of the dataset size) resulted in high AUPR across various datasets. Similarly, we also examined various minimum and maximum windows, and found that our default values resulted in optimal AUPR (**Supplementary Figure 15b**) across datasets of varying sizes. However, users may want to modify these parameters to increase or decrease the leniency of the weighting for a number of reasons, such as when applying Epoch to more complex trajectories with large numbers or more complex state changes (and correspondingly, a large number of epochs). In this case, we would recommend finding optimal lag and window in an analogous method. Specifically, this would entail designing a modular network (and corresponding GRN) representative of the more complex trajectory, using this network to simulate synthetic datasets, and performing a parameter sweep to optimize reconstruction AUPR. Synthetic network design and simulation for both simple and complex trajectories can be done via platforms such as Dyngen.

### Dynamic Network Extraction

Epoch will extract a dynamic network from the reconstructed static network. Specifically, the process begins by breaking pseudotime into “epochs,” or time periods. A number of options are available to users to accomplish this. Briefly, Epoch can define the epochs based on pseudotime, equal cell ordering (resulting in equal number of cells per epoch), k-means or hierarchical clustering, sliding window similarity, or user-defined manual assignment. With the exception of user-defined assignment and sliding window similarity, the number of epochs is specified by the user, and can be determined by examining the heatmap of gene expression across pseudotime and estimating the rough number of expression states represented in the data. We recommend smoothing the data to aid in heatmap visualization, which can be done through the Epoch. Alternatively, it is unnecessary to supply the number of epochs if using the similarity method, which will automatically detect epochs based on correlation between groups of cells along a sliding window across pseudotime. After epochs are defined, Epoch will assign genes to epochs, based on their activity along pseudotime. In brief, this is either based on activity (i.e. genes active in any epochs will be assigned to those epochs), or based on differential expression (i.e. genes are assigned based on if they are differentially expressed in an epoch). Specifically, if genes are assigned by activity, Epoch will compute, for each gene, a threshold against which average expression of the gene in an epoch is compared. This threshold can be modified by user input. If instead genes are assigned by differential expression, a p-value threshold is used to determine assignments. Finally, “orphan genes,” which we define as dynamically expressed genes that do not get assigned to any epoch, are assigned to the epoch in which their average expression is maximum.

Based on these assignments, Epoch will fracture the static network into a dynamic one composed of “epoch networks” and “transition networks”. Specifically, an edge between regulator and target gene appears in an epoch network if the regulator is assigned to that epoch. Further, an edge will appear in a transition subnetwork under two conditions: (1) For an activating edge, the target is not active in the source epoch, is active in the subsequent epoch, and regulator is active or (2) For a repressive edge, the target is active in the source epoch, is not active in the subsequent epoch, and the regulator is active. Following this step, Epoch will return a dynamic network represented by a list of individual GRN tables. This includes the epoch networks or essentially the dynamic network, as well as transition networks that describe how an epoch network may transition into a subsequent epoch network.

### Top Regulator Prediction

Epoch employs two graph theoretic methods to predict “top regulators,” the transcription factors that appear to have the most influence in driving changes in or maintaining topology. First, Epoch will rank regulators by weighted PageRank. In brief, the PageRank centrality measures the importance of nodes in a network based on the number and quality of links of which a node is a part. In essence, the most influential nodes are likely to interact with many influential nodes. Second, Epoch will rank regulators by the product of normalized betweenness and normalized degree. Here, the assumption is that the most influential nodes are likely to be traversed by many shortest paths (and thus have high betweenness) and interact with many other nodes (and thus have high degree). In Epoch, PageRank, betweenness, and degree centralities are implemented through the ‘igraph’ package. By default, cross-weighted edge weights are used, but can be further specified by the user. In both top regulator prediction methods Epoch will return ranked lists of nodes, and further specify their corresponding PageRank or normalized betweenness, normalized degree, and betweenness-degree product.

## Supplementary materials

**Supplementary Figure 1.**
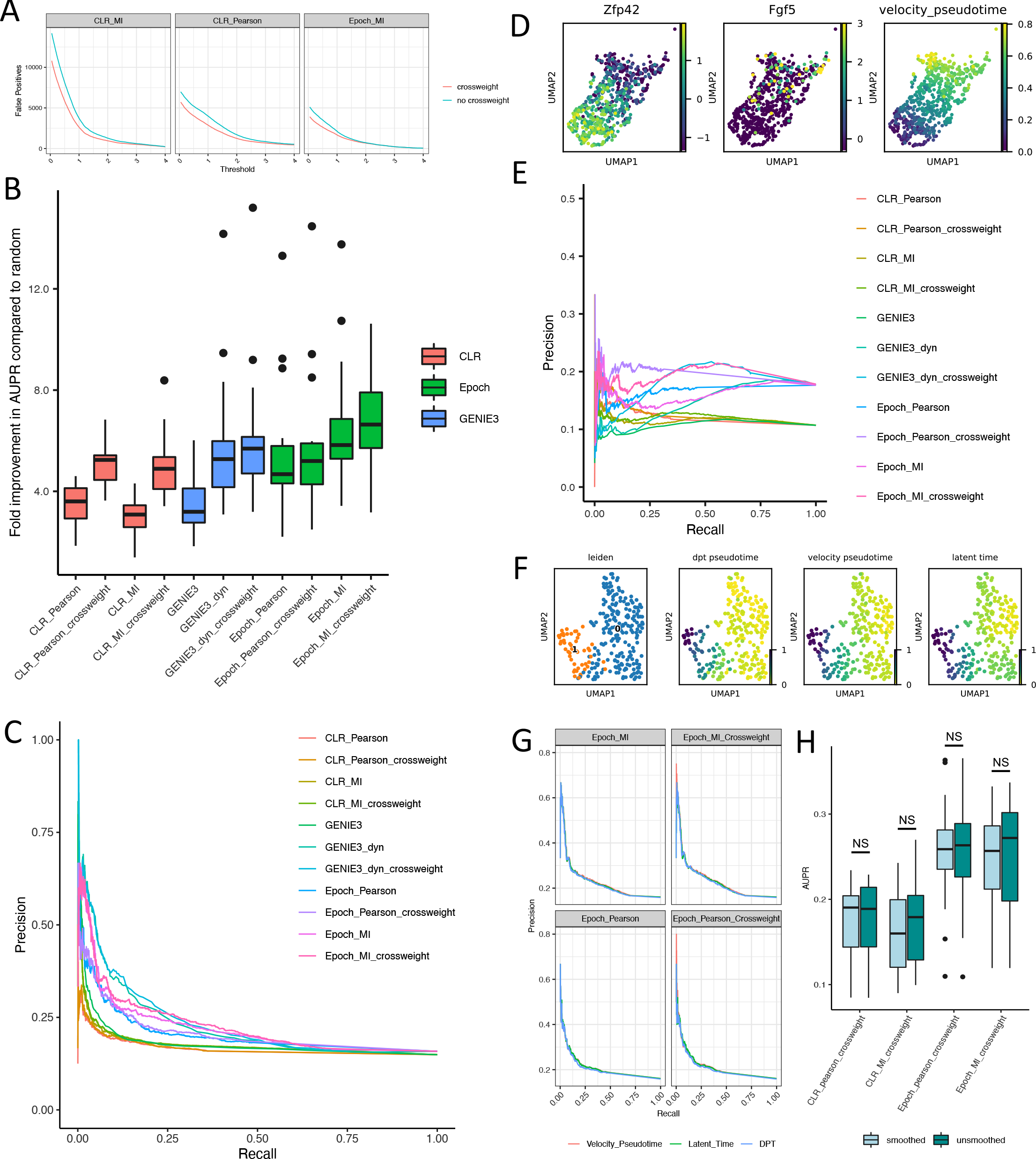
Epoch benchmarking continued. (A) An example from GRN reconstruction on a synthetic dataset showing the difference in number of false positives based on whether or not cross-weighting is performed. (B) Full results of benchmarking reconstruction of synthetic data. Four versions of CLR (Pearson vs. MI, cross-weighting vs. no cross-weighting), four versions of Epoch (same as CLR), and three versions of GENIE3 (original, limited to dynamic genes, and with cross-weighting) are compared. Kruskal-Wallis p=1.75×10^-11^. (C) Full results of benchmarking reconstruction of the muscle development data. Methods are the same as in A. (D) Early mESC directed differentiation data (subsetted from Spangler et al. 2018). Zfp42 expression, Fgf5 expression, and velocity pseudotime are shown. (E) Full results of benchmarking reconstruction of the early Spangler et al. data. Methods are the same as in A. (F) Clustering, diffusion pseudotime, velocity pseudotime, and latent time of the E12.5 muscle development data. (G) Precision recall curves of four variations of Epoch using three different pseudotime annotations. (H) Effect of smoothing vs. unsmoothing on AUPR of reconstruction using various methods.

**Supplementary Figure 2.**
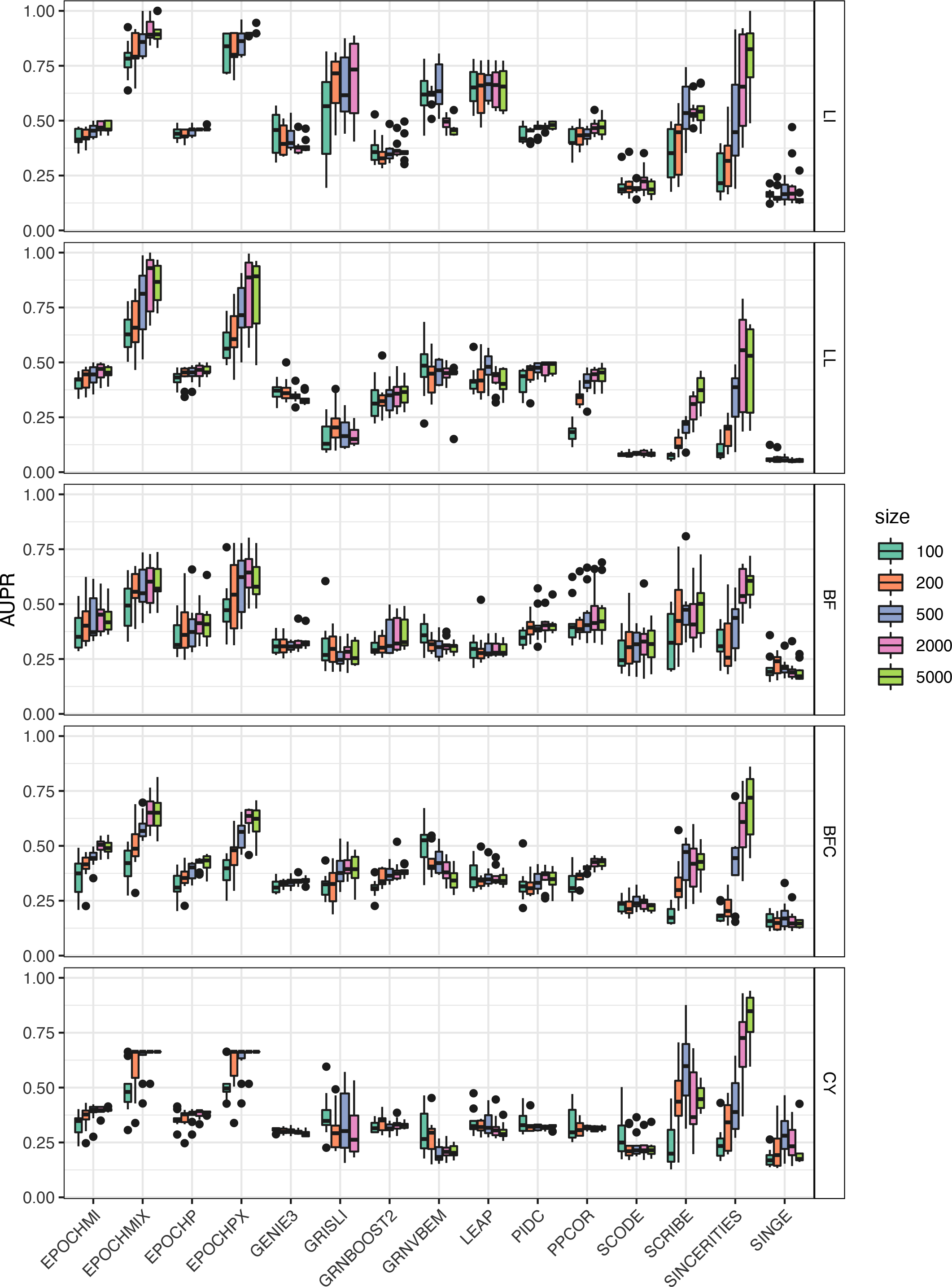
BEELINE synthetic data benchmarking. AUPR of reconstruction on synthetic data using the four versions of Epoch (MI = mutual information, P = Pearson correlation, X = with cross-weighting) and 11 other single-cell GRN reconstruction tools. Results are faceted by trajectory type (LI = linear, LL = long linear, BF = bifurcating, BFC = bifurcating converging, CY = cycle) and colored by number of cells in the dataset.

**Supplementary Figure 3.**
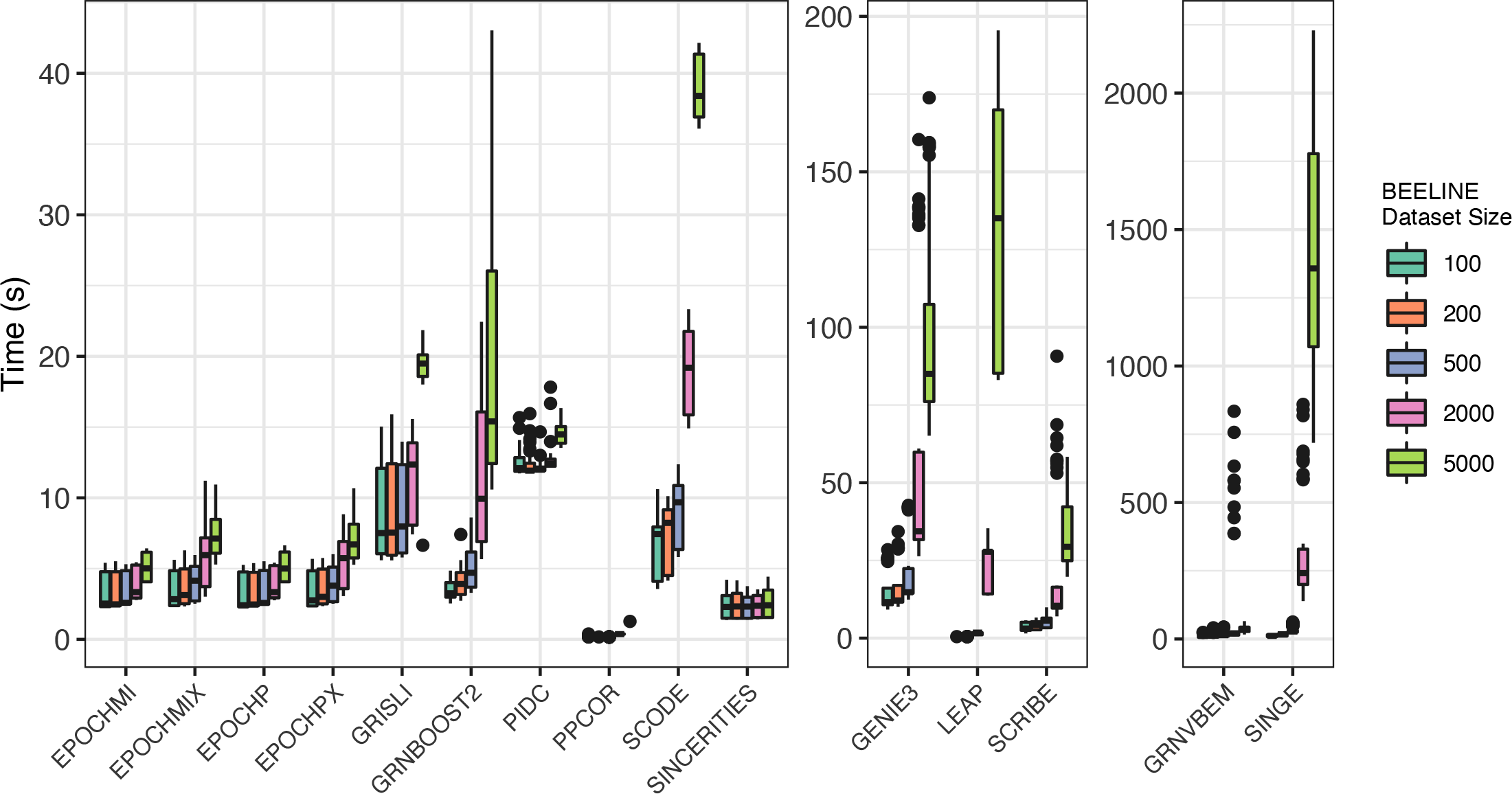
BEELINE synthetic data runtimes. Runtimes (in seconds) are plotted for each method, faceted by magnitude of runtime. Four versions of Epoch are shown (MI = mutual information, P = Pearson correlation, X = with cross-weighting), along with 11 other single-cell GRN reconstruction tools. Data is colored based on number of cells in the dataset.

**Supplementary Figure 4.**
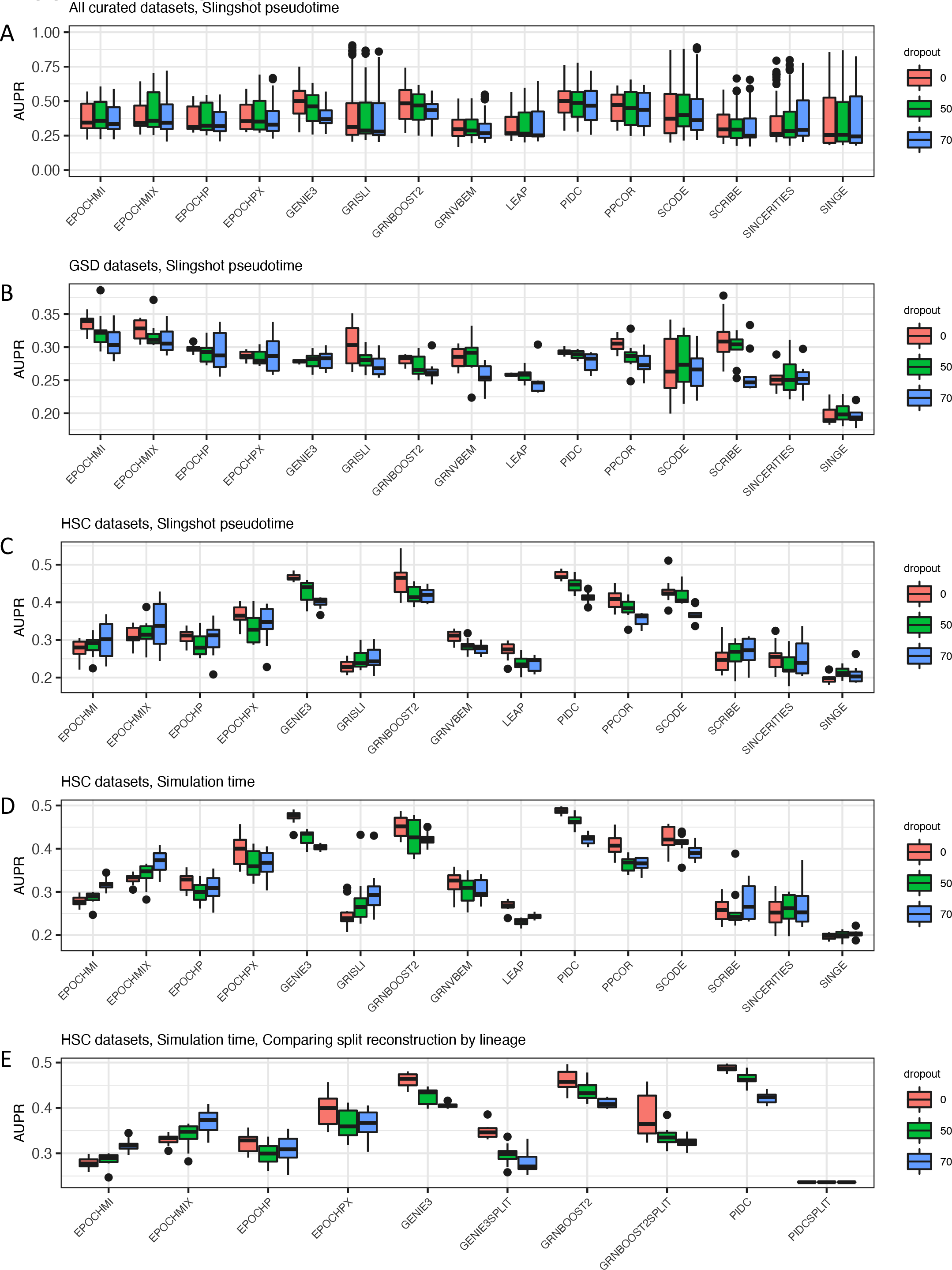
BEELINE curated data benchmarking. AUPR of reconstruction on curated data (as described in Pratapa et al. 2020), using the four versions of Epoch (MI = mutual information, P = Pearson correlation, X = with cross-weighting) and 11 other single-cell GRN reconstruction tools. Panel (A) summarizes the results from all curated datasets. Panel (B) shows results from reconstruction on the GSD datasets using BEELINE-provided Slingshot pseudotime. Panel (C) shows results from reconstruction on the HSC datasets using BEELINE- provided Slingshot pseudotime. Panel (D) shows results from reconstruction on re-simulated HSC datasets using true simulation time from BoolODE (Pratapa et al., 2020). Panel (E) shows results from reconstruction on re-simulated HSC datasets using true simulation time, comparing the results of reconstructing using GENIE3, GRNBoost2, and PIDC with no splitting vs. splitting along each lineage before compiling (SPLIT = with splitting and compiling).

**Supplementary Figure 5.**
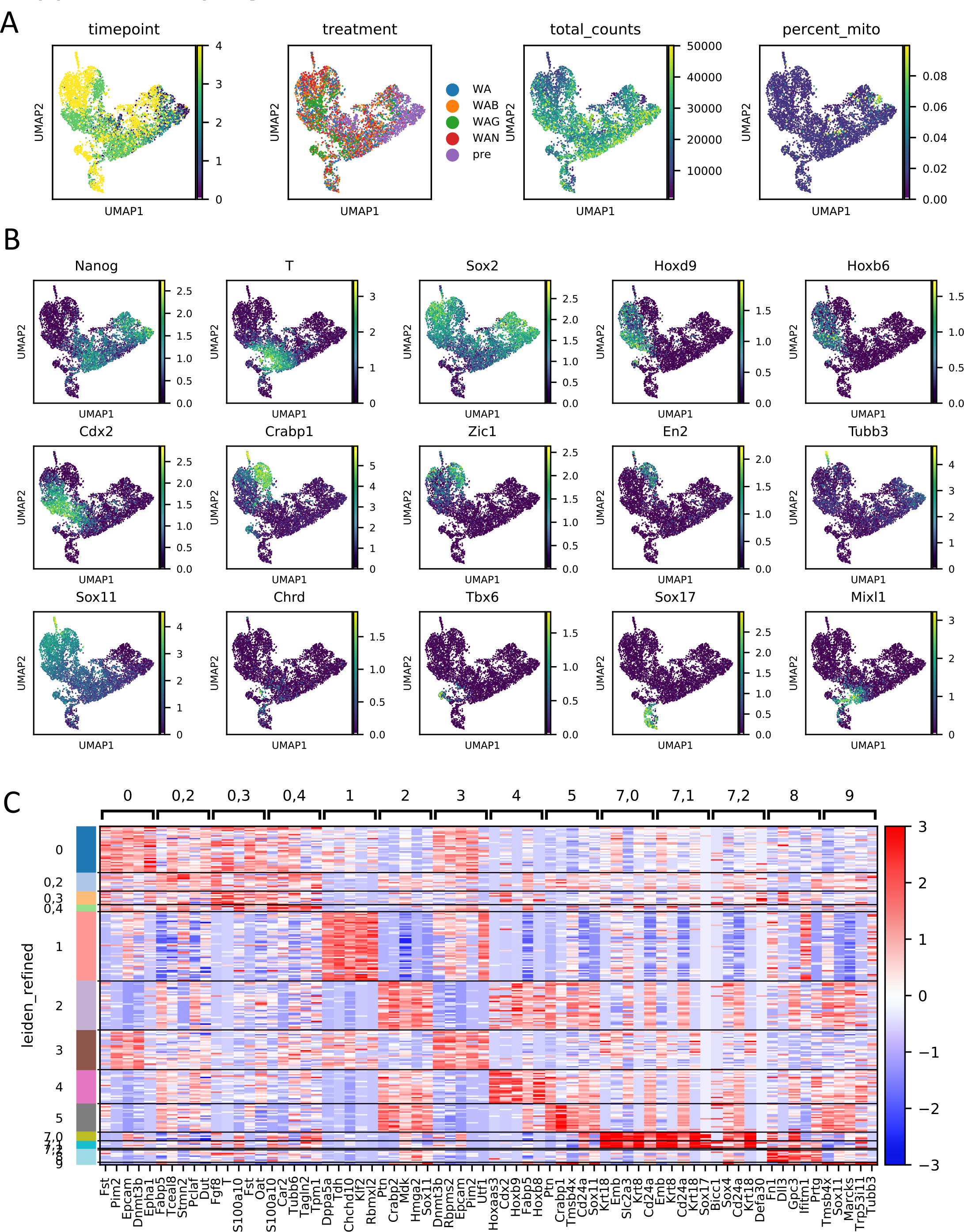
MULTI-seq directed differentiation cell typing. (A) timepoint, treatment, total counts, and percent mitochondrial transcripts content of the MULTI-seq data. (B) Select marker gene expression of the MULTI-seq data. (C) Differential expression analysis of the MULTI-seq data by cluster.

**Supplementary Figure 6.**
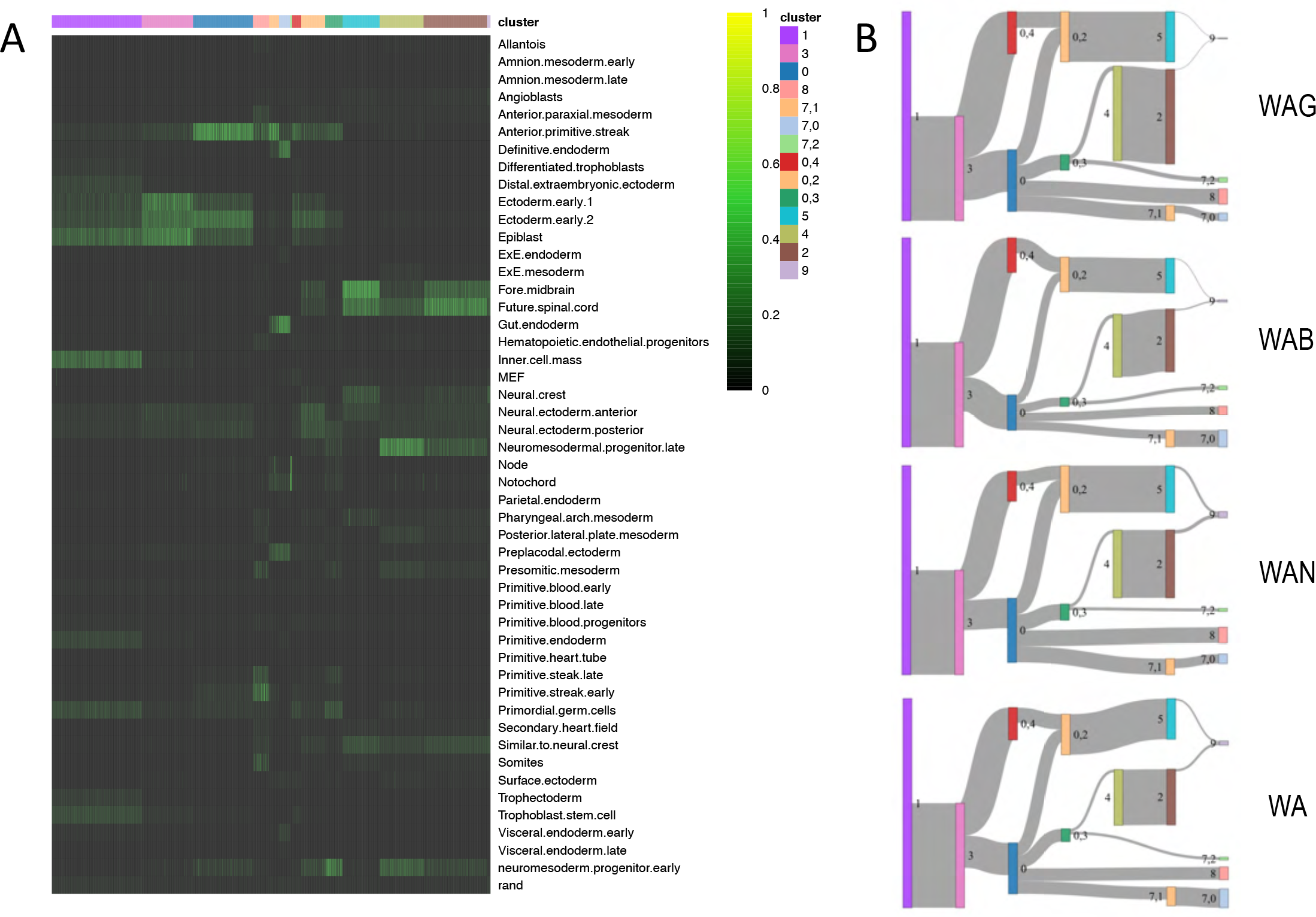
MULTI-seq cell type classification and cluster transitions. (A) SingleCellNet classification results. The classifier was trained from mouse embryo data found in literature, curated by members in the lab. (B) Sankey diagrams depicting cluster transitions (and cell fate transition proportions) as broken down by treatment.

**Supplementary Figure 7.**
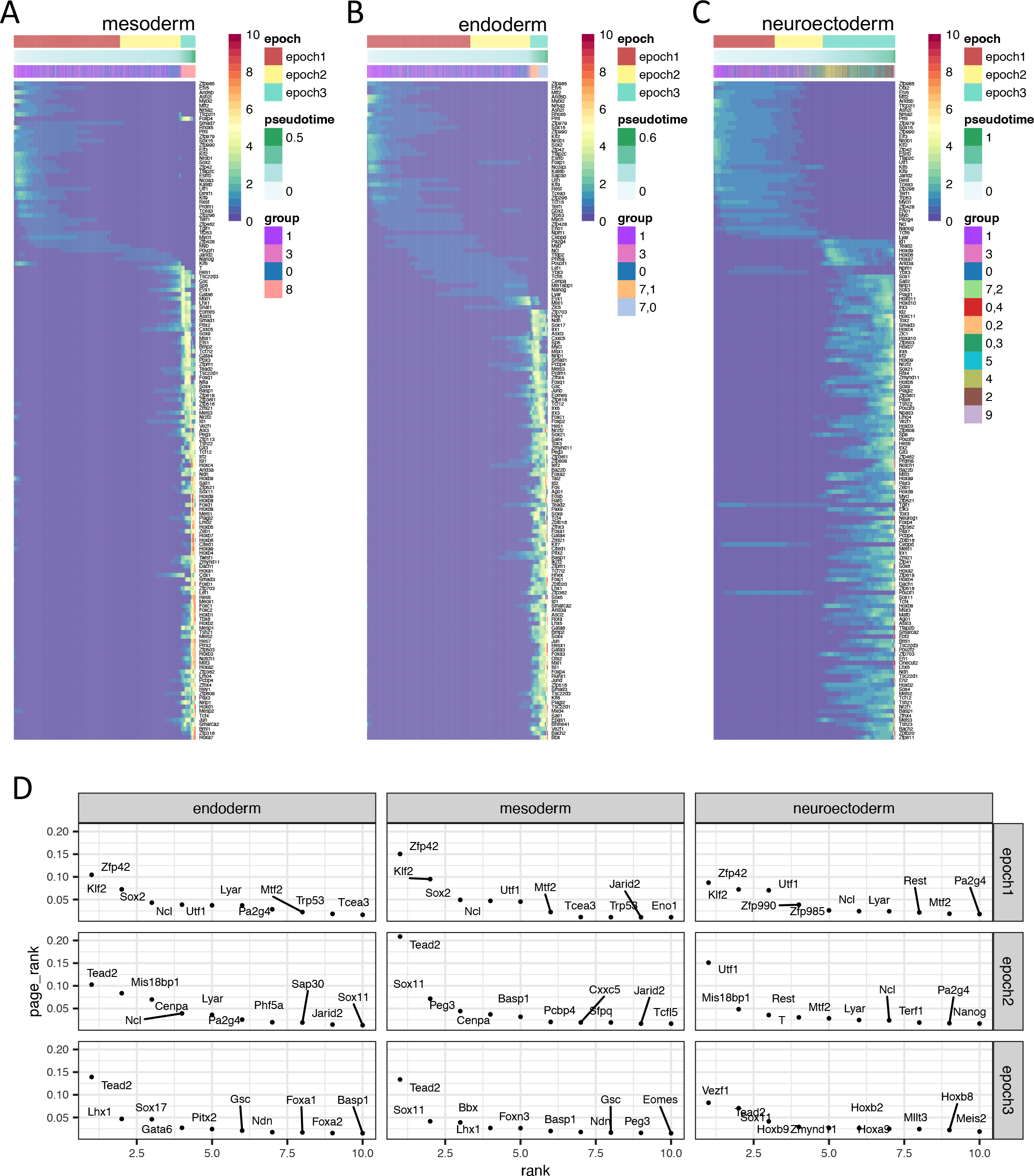
MULTI-seq network reconstruction and top regulators. Heatmaps depicting TF expression across time and Epoch definitions along the (A) mesoderm path, (B) endoderm path, and (C) neuroectoderm path. (D) Top 10 regulators in each epoch network along each germ layer path as predicted by Epoch via PageRank.

**Supplementary Figure 8.**
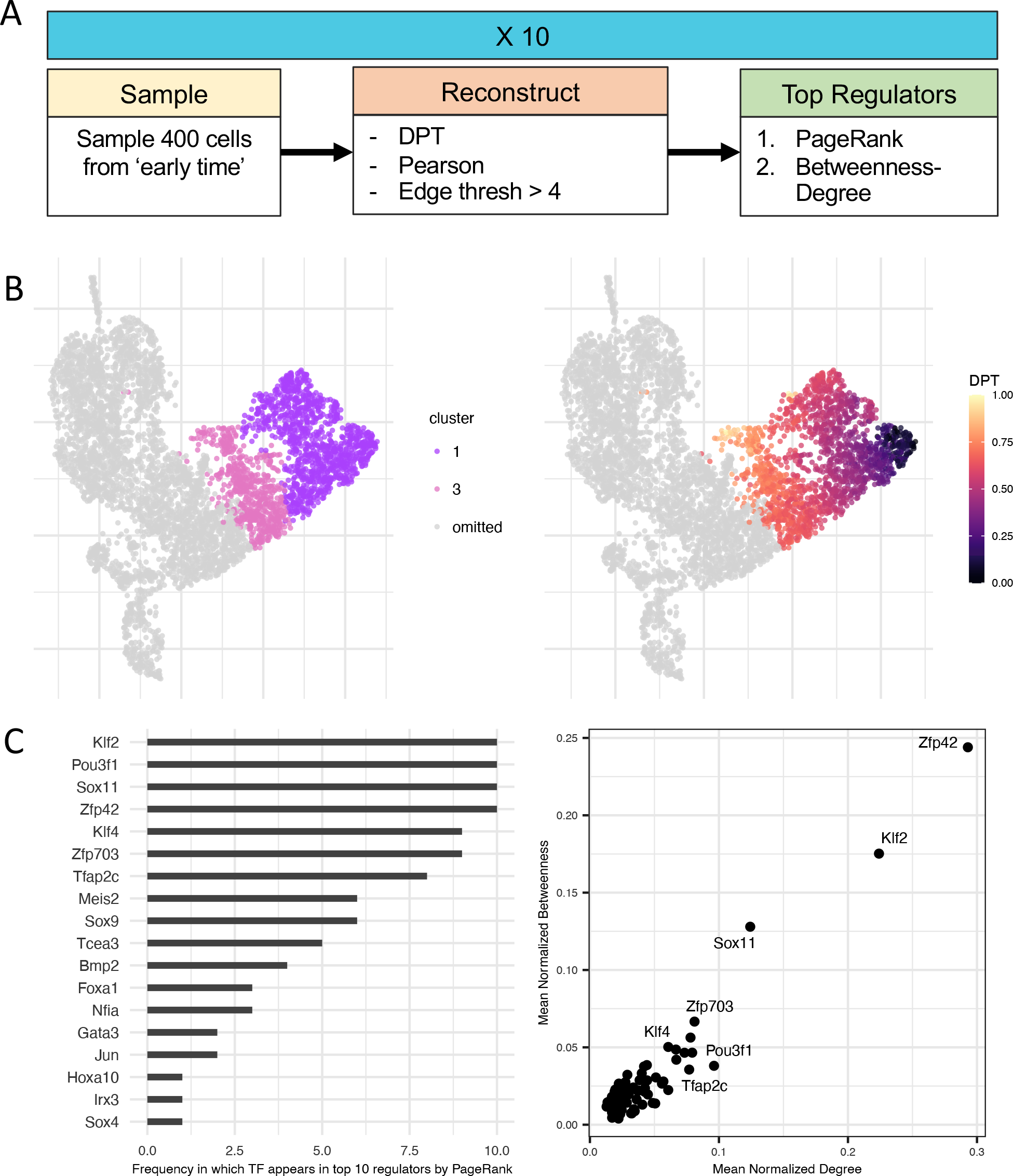
Bootstrapped top regulator prediction for inner cell mass to epiblast-like transition. (A) Workflow for bootstrapping network reconstruction in order to predict top regulators. (B) Only cells from early time were used. Clusters used and corresponding diffusion pseudotime shown here. (C) Top regulators for this early fate transition as predicted using PageRank and Betweenness-Degree.

**Supplementary Figure 9.**
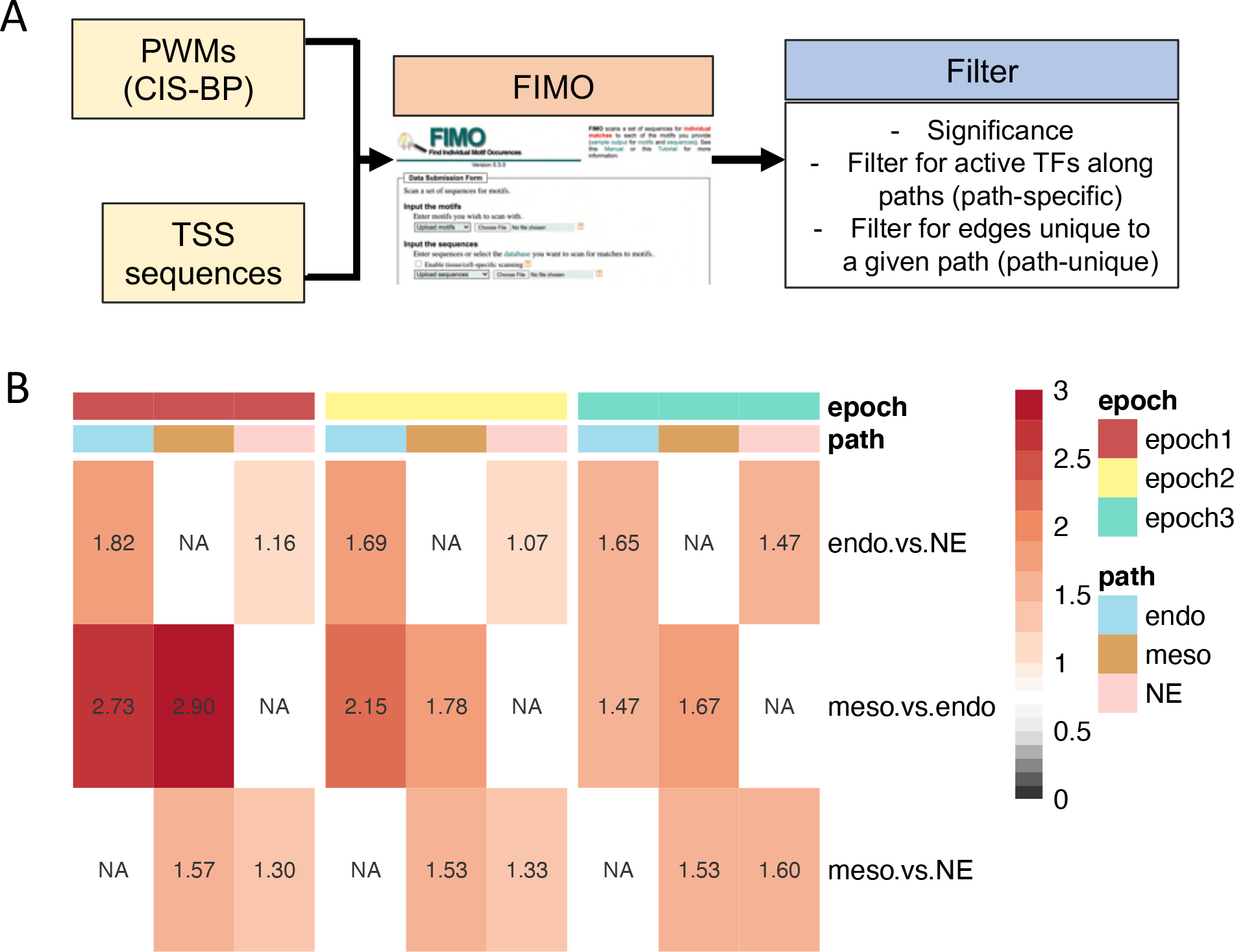
Proof of concept for differential network analysis. (A) Path-unique gold standard creation. (B) Performance (AUPR) over random of differential network in recovering path-unique gold standard edges. Each row is a comparison between two paths. Each column represents unique edges recovered for a given path. As an example, the top left box indicates that in the endoderm vs. neuroectoderm comparison, in the first epoch the endoderm differential network is 1.82 times better than a shuffled network in recovering edges that are specific to endoderm (and not present in neuroectoderm).

**Supplementary Figure 10.**
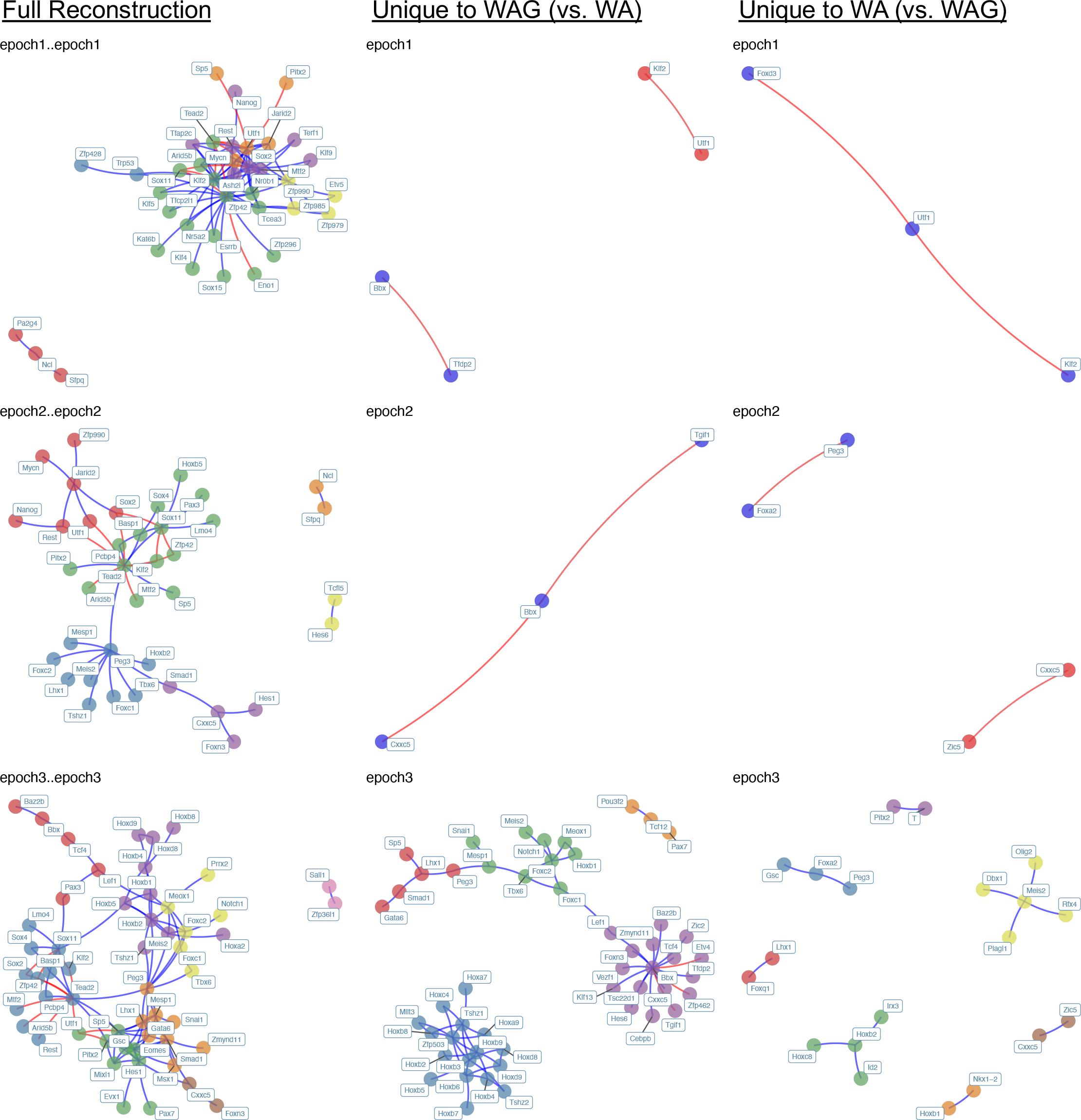
Mesodermal network and differential networks. The left column (“Full Reconstruction”) depicts the dynamic mesodermal network. The middle and right columns depict the WAG vs. WA differential network. Middle is the network unique to WAG, right is the network unique to WA. TFs are colored by community. Blue and red edges represent activating and repressive edges respectively.

**Supplementary Figure 11.**
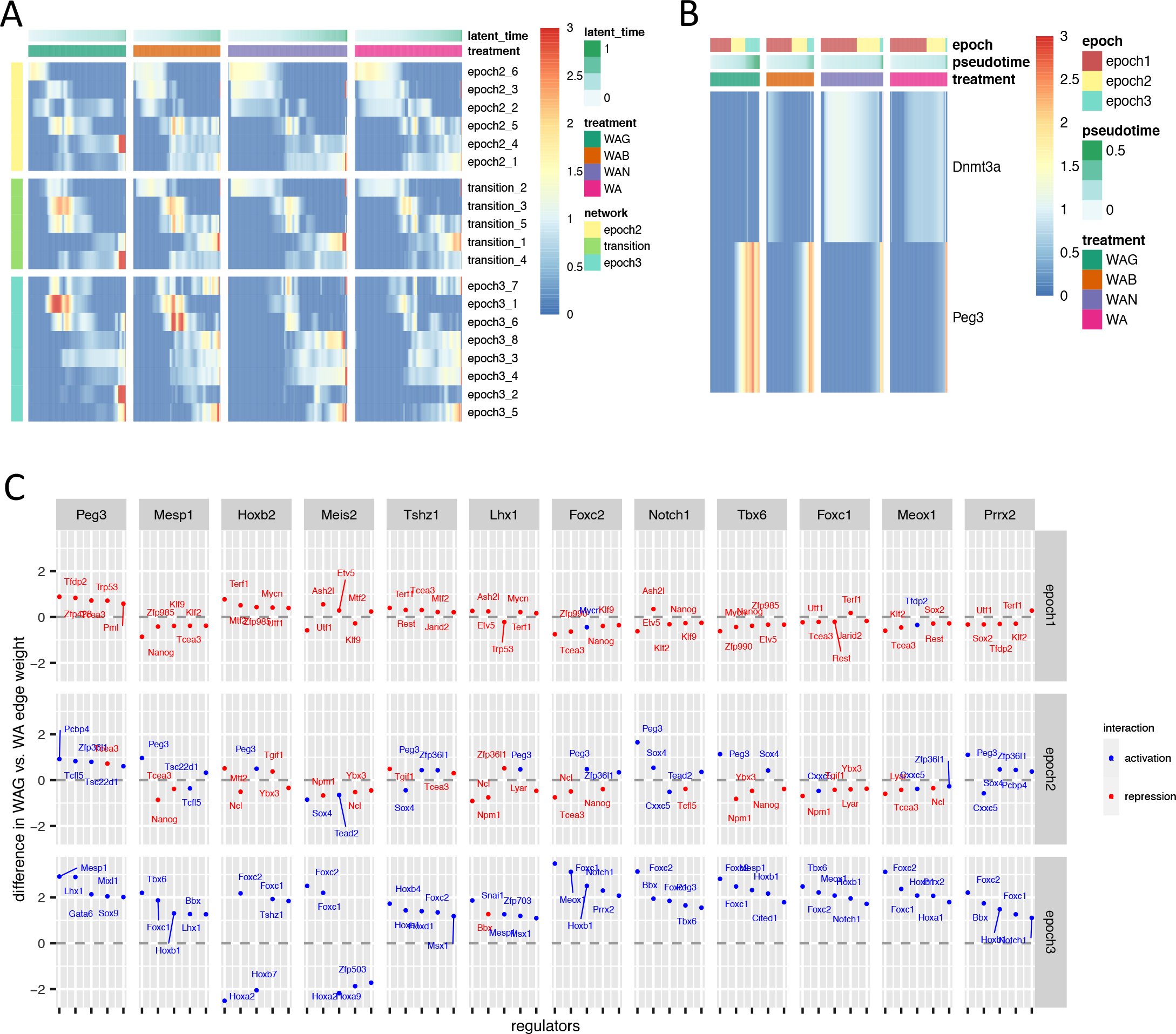
Mesodermal network analysis continued. (A) Average community expression over time by treatment along the full dataset. Communities shown (each row) are from the epoch 2 subnetwork (yellow), transition (green), and epoch 3 subnetworks (aqua) of the mesodermal network. (B) Peg3 and Dnmt3a expression across time along the mesodermal path of the MULTI-seq data. (C) Top differential regulators of mesodermal genes. Blue and red represent activators and repressors respectively. Y-axis is plotting difference in edge weights between the WAG and WA networks. Thus, TFs appearing above the 0-line are more unique to the WAG network, and those below are more unique to the WA network.

**Supplementary Figure 12.**
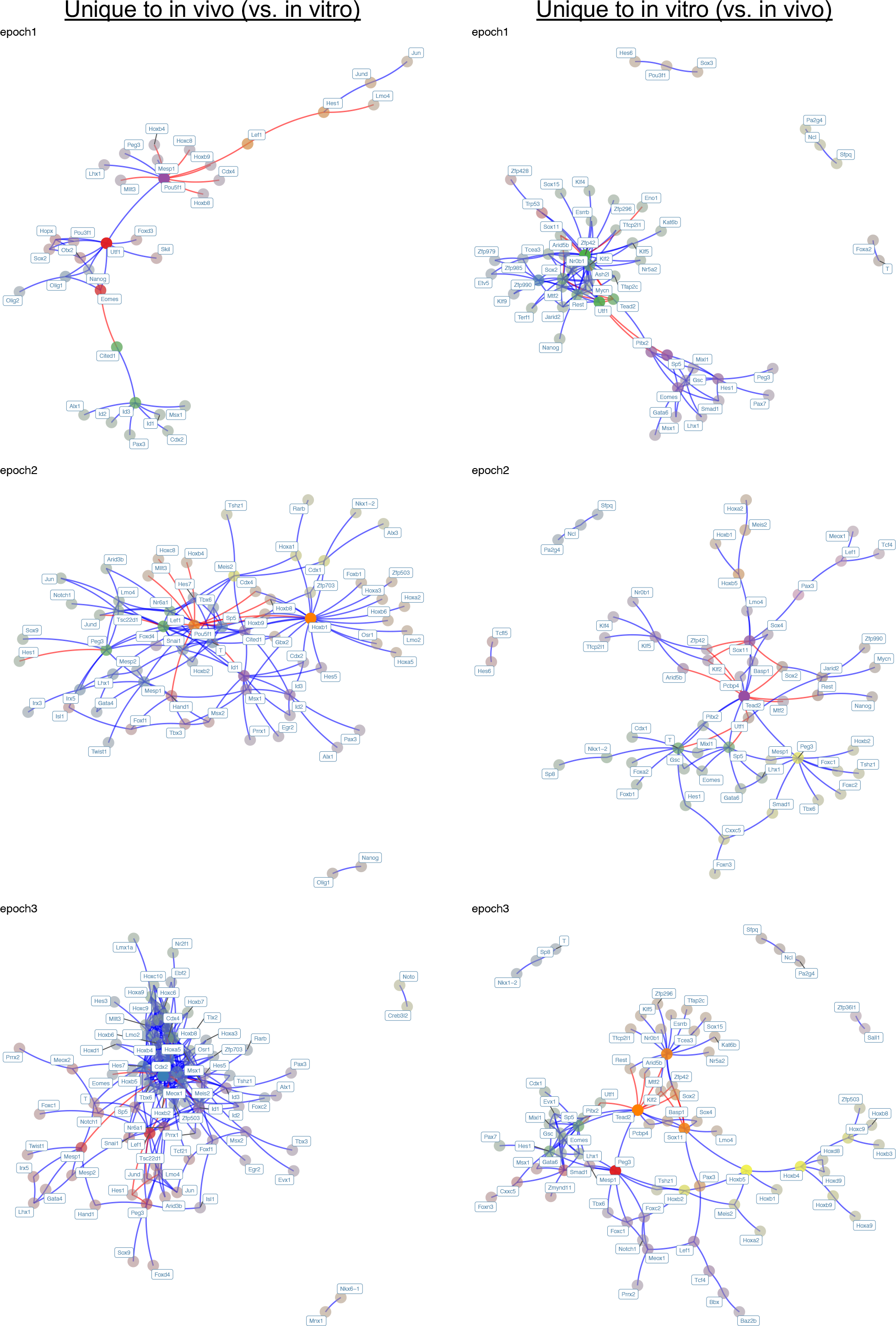
*In vivo* vs. *in vitro* mesodermal differential network. (Left) The differential network containing edges unique to the *in vivo* network. (Right) The differential network containing edges unique to the *in vitro* network.

**Supplementary Figure 13.**
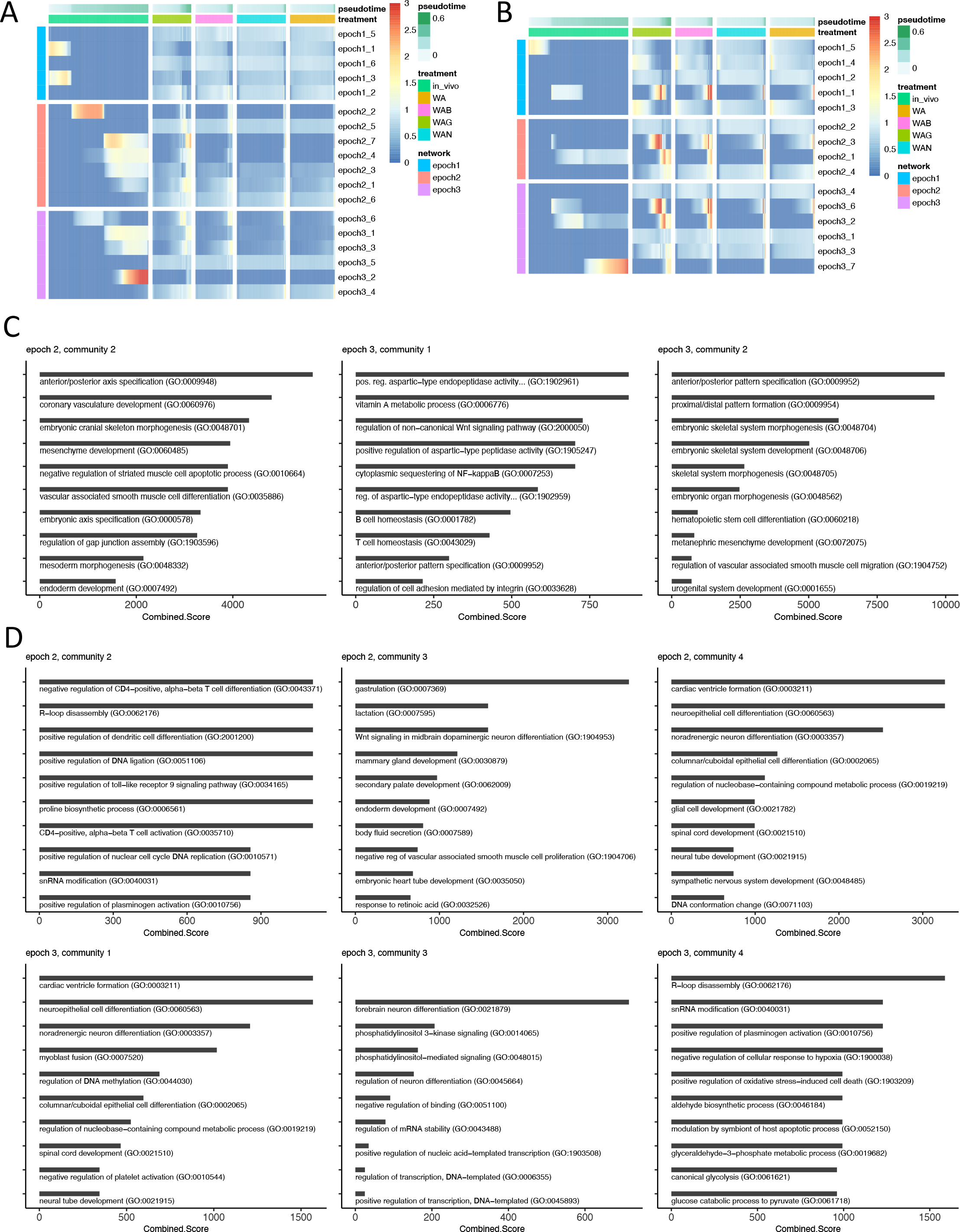
*In vivo* vs. *in vitro* mesodermal network comparison continued. (A) Average *in vivo* community expression over time along the mesodermal path for the *in vivo*, *in vitro* WAG, *in vitro* WAB, *in vitro* WAN, and *in vitro* WA data. (B) Average *in vitro* community expression over time along the mesodermal path for the *in vivo*, *in vitro* WAG, *in vitro* WAB, *in vitro* WAN, and *in vitro* WA data. (C) Top ten enriched terms for *in vivo*-specific modules based on the Combined Score from Enrichr GSEA analysis. The three modules included are activated in the *in vivo* data but not (or weakly activated) in the *in vitro* data (see panel A). (D) Top ten enriched terms for *in vitro*-specific modules based on the Combined Score from Enrichr GSEA analysis. The six modules included are activated in the *in vitro* data but not (or weakly activated) in the *in vivo* data (see panel B).

**Supplementary Figure 14.**
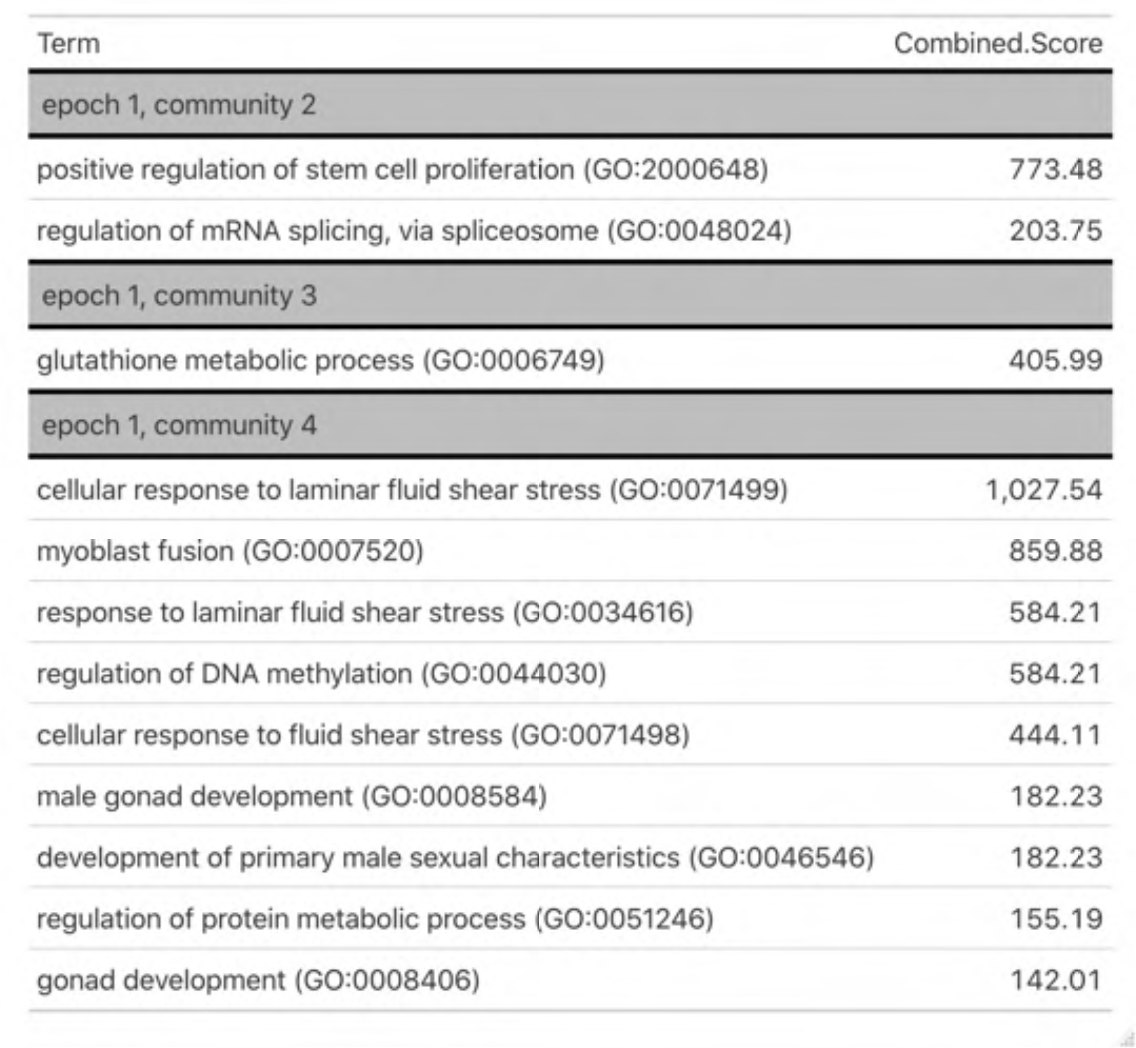
Early functional differences in *in vitro* vs. *in vivo* networks. Enriched terms for *in vitro*-specific modules based on the Combined Score from Enrichr GSEA analysis, specific to the first epoch. These three communities belong to the first epoch and are activated in the *in vitro* data but not (or weakly activated) in the *in vivo* data (see Supplementary Figure 3b).

**Supplementary Figure 15.**
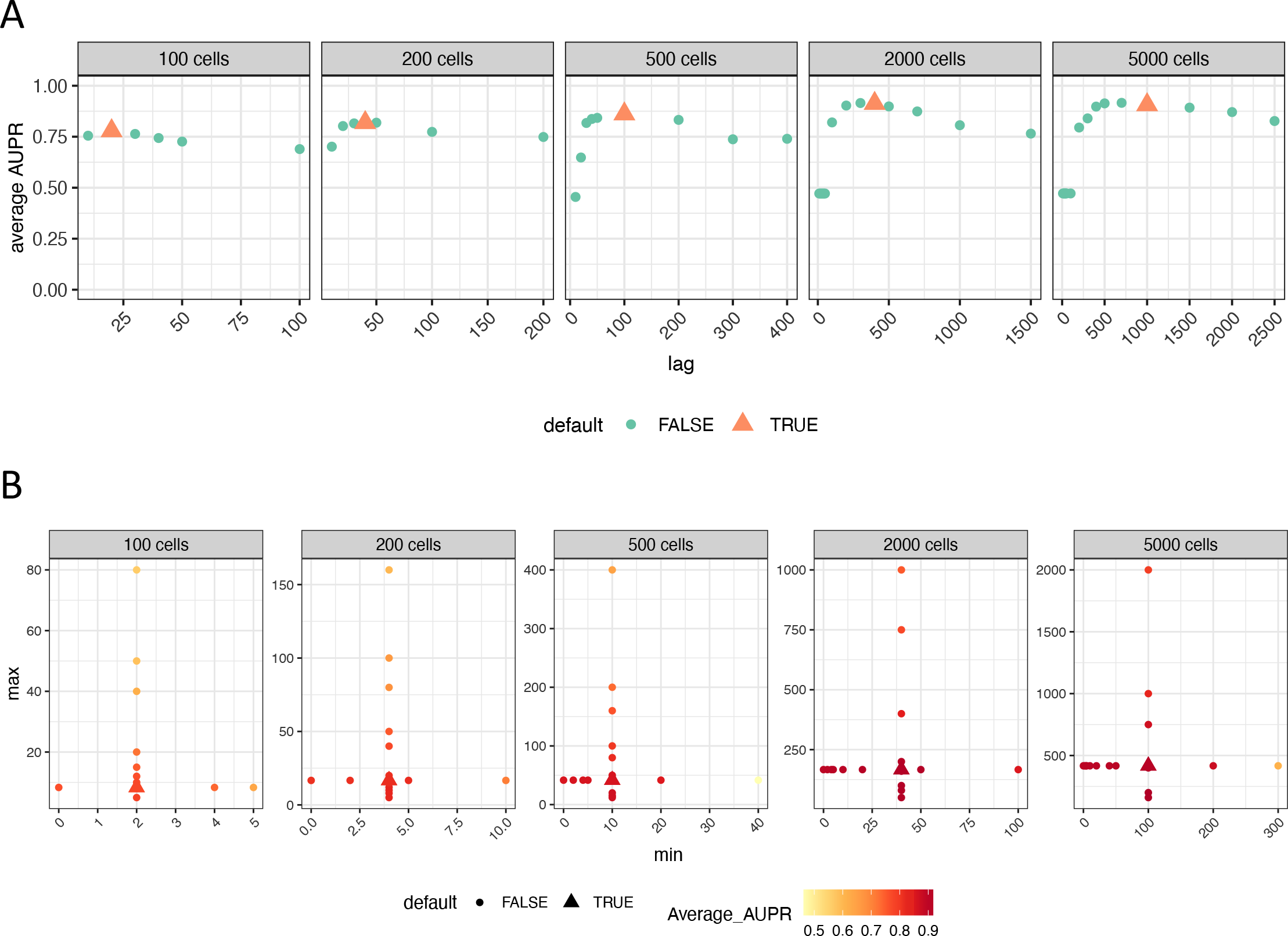
Optimal default parameter settings for synthetic linear trajectories. (A) Average AUPR of reconstruction using different cross-weighting lag times across datasets of varying sizes, taken from BEELINE linear datasets. Each dataset size contains 10 distinct datasets, Default lag time is plotted as the orange triangle, and is computed as one-fifth of the number of cells in the data. (B) Average AUPR of reconstruction using different minimum and maximum windows in cross-weighting. Data shown is reconstruction from BEELINE linear datasets. Each dataset size contains 10 distinct datasets. Default minimum and maximum is indicated by the triangle, and is defined as 1/50 and 1/12 of total cells respectively. High average AUPR is indicated by the darker red colors.

**Supplementary Table 1.** Cell type annotation by cluster.

| Cluster | Cell Type Label | Evidence |
| --- | --- | --- |
| 1 | Undifferentiated ESCs | This cluster classifies strongly as inner cell mass and epiblast. It expresses a number of pluripotency genes including Zfp42. |
| 3 | Epiblast | This cluster classifies as epiblast with some early ectodermal signature. Fgf5 expression begins here. |
| 0 | Primitive Streak | This cluster classifies strongly as primitive streak with some early ectodermal signature. It expresses brachyury. |
| 8 | (mesoderm)<br>Presomitic Mesoderm/Somites | This cluster classifies as presomitic mesoderm and somites, with some residual primitive streak. It expresses a number of mesodermal markers including Mesp1. |
| 7,1 | (endoderm) | 7,0 exhibits strong classification for definitive endoderm and gut endoderm. It expresses definitive endodermal markers including Foxa2 and Sox17. RNA velocity suggests 7,1 cells transition into 7,0, and largely classifies as anterior primitive streak and endoderm. |
| 7,0 |  |  |
| 7,2 | Node | This cluster strongly classifies as node, and expresses Chordin. |
| 0,4 | (neuroectoderm)<br>Early Ectoderm | This cluster classifies weakly as a number of ectodermal cell types, but based on RNA velocity seems to be most similar to early ectodermal cells. |
| 0,2 | (neuroectoderm)<br>Early Ectoderm, Forebrain/ Midbrain fated | This cluster classifies weakly as a number of ectodermal cell types including early ectoderm. Based on RNA velocity, this cluster transitions into cluster 5 (forebrain/midbrain fated). |
| 5 | (neuroectoderm)<br>Future Brain, Forebrain/ Midbrain | This cluster classifies strongly as forebrain/midbrain with some weaker future spinal cord classification. It expresses Zic1 and En2. |
| 9 | (neuroectoderm)<br>Future brain | This cluster classifies weakly as future brain, future spinal cord, and neural crest. It expresses Crabp1 and Tubb3, and RNA velocity suggests it transitions from clusters 2 or 5. |
| 0,3 | Early Neuromesodermal Progenitor | This cluster classifies weakly as a number of cell types including primitive streak and early ectoderm, but more strongly as early neuromesodermal progenitor. RNA velocity shows cells in this cluster transition into a neuromesodermal progenitor. Cells in this cluster express T and some Sox2. |
| 4 | Neuromesodermal Progenitor | This cluster classifies strongly as neuromesodermal progenitor (late and early). It has some T and Sox2 expression. |
| 2 | (neuroectoderm)<br>Future Spinal Cord | This cluster classifies mostly as future spinal cord, and weakly as fore/midbrain. It also expresses many patterning genes from the Hox family indicative of spinal cord fated cells such as Hoxb9. |

